# *NAT10* supports CAG repeat-associated mRNA stabilization and is associated with hepatic metabolic reprogramming during spaceflight

**DOI:** 10.64898/2026.09.17.752365

**Authors:** Ryota Kurimoto, Tomoaki Tanaka, Izumi Ohno, Tomoki Chiba, Takahide Matsushima, Yutaro Uchida, Yuki Naito, Hiroshi Asahara

## Abstract

The regulatory factors underlying posttranscriptional adaptation to spaceflight are incompletely understood. We identified *NAT10*, an RNA acetyltransferase, among RNA modification enzymes significantly upregulated in spaceflight-exposed mouse liver. Transcriptome-wide correlation analysis linked *NAT10* expression to an RNA-metabolism module enriched for CAG repeat density in coding sequences. Actinomycin D chase assays showed that *NAT10* knockdown destabilized CAG repeat-containing transcripts, with *DDX17* serving as the primary validated target; *DCP1A* showed a directionally consistent pattern. Wild-type *NAT10* re-expression fully rescued *DDX17* mRNA stability, the acetyltransferase-dead G641E mutant rescued mRNA stability to a comparable extent, and the K290A helicase-domain mutant failed to rescue, consistent with a requirement for helicase domain integrity, whereas acetyltransferase activity was not strictly required under these experimental conditions. RNA immunoprecipitation further supported preferential *NAT10* association with CAG repeat-containing mRNAs, with reduced association after single-site CAGCAG deletion in *DDX17* and *DCP1A* reporters. Gene set enrichment analysis associated *NAT10* upregulation in spaceflight liver with suppressed fatty acid beta-oxidation and induced cholesterol biosynthesis; *NAT10* depletion in hepatocellular carcinoma cells selectively opposed the cholesterol-associated, but not the broader metabolic, component of this signature. These findings support a model linking *NAT10* to CAG repeat-associated mRNA stabilization in the context of hepatic metabolic and posttranscriptional to spaceflight.

## Introduction

Long-term spaceflight introduces a combination of extreme environmental challenges, including microgravity, cosmic radiation, disrupted circadian rhythms, and confinement-related stress. These stressors disrupt homeostatic systems in nearly every organ, leading to alterations in immune responses, cardiovascular physiology, and metabolic homeostasis ^1,2^. Microgravity has been reported to stimulate autophagy and proteasomal activity in hepatic tissue and increase oxidative stress ^3^. Liver injury, inflammation, and alterations in carbohydrate and lipid metabolism have been observed^4^.

Although the transcriptional adaptation to spaceflight has been studied extensively, the involvement of RNA modifications in organismal responses to spaceflight is largely uncharacterized. Growing evidence implicates epitranscriptomic regulation—the chemical modification layer of RNA—as a pivotal determinant of cellular adaptation under stress. The repertoire of characterized RNA modifications now exceeds 150 chemically distinct species, including RNA marks such as N6-methyladenosine (m6A), 5-methylcytosine (m5C), and pseudouridine (Ψ), as well as emerging marks including N4-acetylcytidine (ac4C) ^5–10^. These modifications play key roles in mRNA processing, translation, and decay, influencing gene expression during cellular stress responses, differentiation, and development ^5,10^. Advanced sequencing techniques have enabled transcriptome-wide mapping of these modifications, which indicates their dynamic nature and regulatory functions ^6,9^.

Despite advances in understanding individual RNA modifications, whether the broader landscape of RNA modification enzymes responds to the unique stressors of spaceflight is largely unknown. Spaceflight imposes a distinct combination of microgravity, cosmic radiation, and disrupted circadian cycles, which may engage posttranscriptional regulatory mechanisms not observed in conventional in vitro stress models. Therefore, systematic characterization of RNA modification enzyme expression across spaceflight-exposed tissues could identify novel epitranscriptomic regulators and provide mechanistic insights into transcriptome-wide alterations associated with the space environment.

*NAT10* (N-acetyltransferase 10) has been identified as the sole acetyltransferase that deposits ac4C onto RNA ^11,12^. *NAT10* also functions as a nucleolar protein that is involved in ribosome biogenesis and nuclear architecture ^13,14^. *NAT10* catalyzes ac4C formation in rRNA and tRNA in yeast and human cells ^15^. Arango et al. ^11^ demonstrated that ac4C is also present on mRNA, where it promotes transcript stability and translational efficiency; however, subsequent studies have highlighted the uncertainty regarding the presence and functional significance of mRNA acetylation ^16,17^.

NAT10-mediated ac4C deposition has been implicated in diverse biological contexts, including oncogenesis, cellular differentiation, and immune regulation ^18,19^. Substantial work has characterized the involvement of *NAT10* in disease, encompassing both ac4C-dependent and ac4C-independent mechanisms ^20^.

However, little is known about the behavior of *NAT10* in extreme environments such as spaceflight, or whether it functions as part of a conserved cellular adaptation program. Furthermore, the means through which *NAT10* participates in broader RNA degradation pathways or whether it acts in concert with motif-driven transcript selection is unclear.

Although AU-rich elements are well-established determinants of mRNA stability, with UUAUUUAUU identified as a key destabilizing motif ^21^, some motifs are underexplored in the context of stress-responsive RNA regulation. CAG repeat motifs in CDS, which contribute to neurodegenerative conditions, including Huntington’s disease, are associated with R-loop formation, repeat instability, translational effects, RNA export, and RNA decay ^22–27^; however, the role of CAG repeat motifs in stress-responsive RNA regulation is uncharacterized.

Here, we conducted a systematic survey of RNA modification enzyme expression across spaceflight-exposed mouse tissues, leveraging openly deposited RNA-seq data from the NASA GeneLab Data Repository (GLDS-168, GLDS-420, GLDS-599, GLDS-419, GLDS-397, and GLDS-102). Among the 152 RNA modification-related genes examined, *NAT10* was among a set of RNA modification enzymes substantially upregulated in the livers of spaceflight-exposed mice, distinguished by statistically significant upregulation in spaceflight liver. Subsequent transcriptomic correlation analyses, motif-based gene enrichment analyses, and functional assays indicated that *NAT10* expression was associated with a module of RNA degradation factors and transcripts enriched for CAG repeat density. Actinomycin D-based mRNA stability assays and RNA immunoprecipitation experiments further supported a model in which *NAT10* preferentially stabilizes selected CAG repeat-containing mRNAs through a mechanism involving RNA association and helicase-domain integrity. Together, these findings identify *NAT10* as a candidate posttranscriptional regulator associated with the spaceflight response, with potential roles in mRNA stabilization and hepatic metabolic gene regulation.

## Results

### Systematic screening of RNA modification enzymes identifies *NAT10* as a spaceflight-responsive regulator in mouse liver

To characterize the transcriptomic response to spaceflight and identify putative epitranscriptomic regulators, we analyzed RNA-seq data from spaceflight-exposed (FLT, n = 5) and ground control (GC, n = 5) mouse liver samples from NASA GeneLab dataset GLDS-168, reprocessed from raw sequencing reads through a STAR–featureCounts–DESeq2 pipeline. A volcano plot of the liver transcriptome indicated a considerable number of transcriptomic alterations (779 upregulated and 1,103 downregulated genes at padj < 0.05 and |logLFC| > 0.5), with *NAT10* among the significantly upregulated genes (logLFC = 0.50, padj = 0.0023; Fig. 1A). To determine whether RNA modification enzyme expression was altered during spaceflight, we examined a curated panel of 152 RNA modification-related genes spanning diverse modification types, including ac4C, m6A, m5C, m1A, m7G, pseudouridine, adenosine-to-inosine editing, and further tRNA and rRNA modification classes. *NAT10* was among a subset of RNA modification enzymes substantially upregulated in spaceflight liver, together with several tRNA-modification writers that showed comparable or larger fold changes (e.g., OSGEPL1/t6A, TARBP1/2’-O-methylation, THUMPD2/m2G; Fig. 1B; Supplementary Fig. 5B), consistent with broader remodeling of the RNA modification enzyme landscape during spaceflight (Supplementary Fig. 5A). Increased *NAT10* expression was confirmed by direct comparison of expression levels across FLT and GC samples (Fig. 1C), and, although a numerically larger fold-change was observed in skeletal muscle, liver was the only tissue among the six examined for which this induction reached statistical significance (Wilcoxon rank-sum p = 0.016; Supplementary Fig. 1). Given this significant upregulation in spaceflight liver and its established role as the sole known ac4C-depositing enzyme in mammalian cells, we selected *NAT10* for downstream functional characterization.

**Figure 1.**
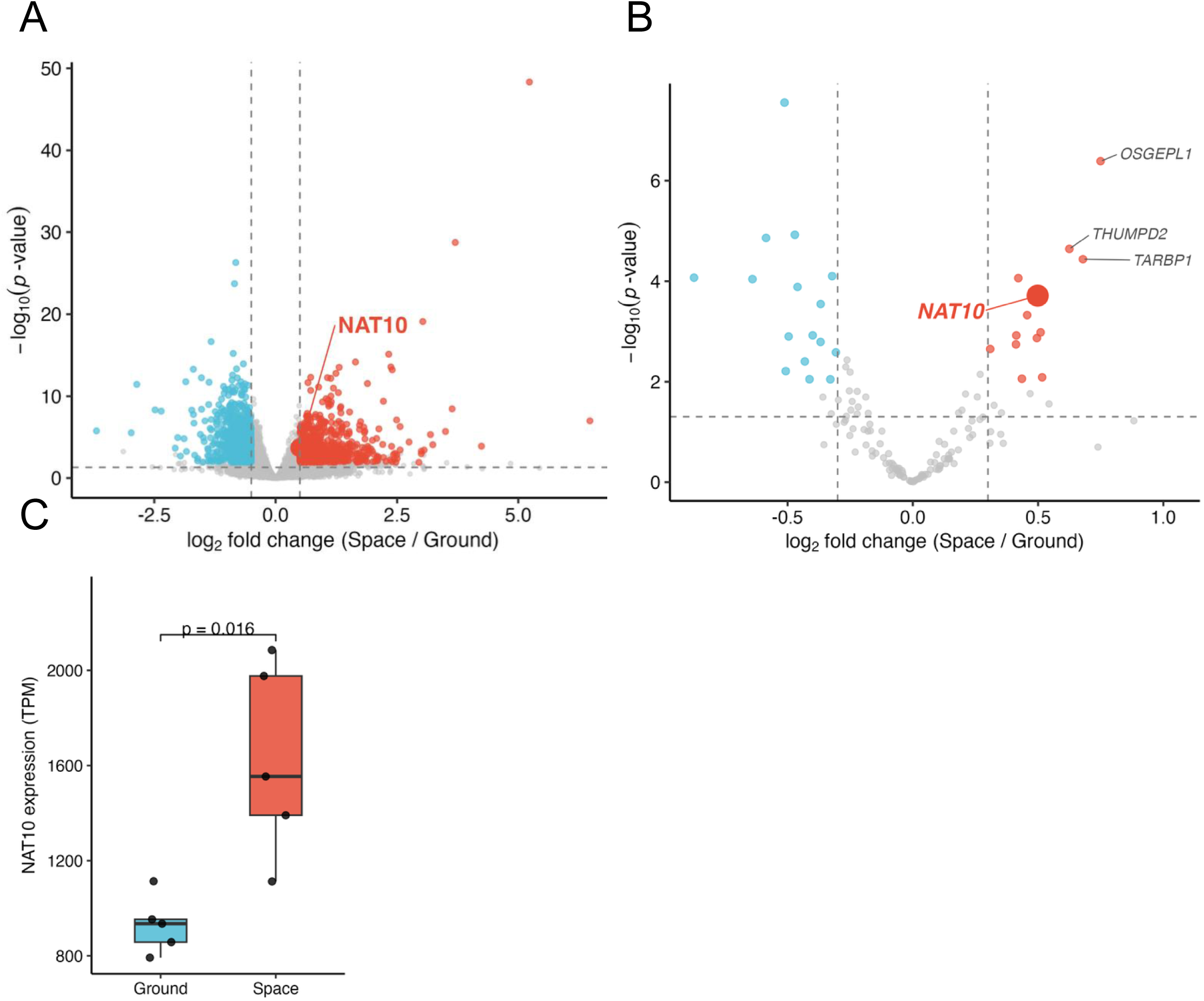
Systematic screening of RNA modification enzymes identifies *NAT10* as a spaceflight-responsive regulator in mouse liver. (A) Volcano plot of all expressed genes in the liver (Space; FLT, n = 5) and ground control (Ground; GC, n = 5) mice from NASA GeneLab dataset (GLDS-168), reprocessed from raw sequencing reads (STAR alignment, featureCounts quantification, DESeq2 differential expression). Each point represents one gene; logL fold change (x-axis, DESeq2 Wald test) plotted against −logLL(p-value) (y-axis). Red: upregulated (padj < 0.05 with logLFC > 0.5); teal: downregulated (padj < 0.05 with logLFC < −0.5); gray: not significant. *NAT10* is highlighted. (B) Volcano plot restricted to the 152-gene RNA modification enzyme panel (DESeq2 results as in A; padj < 0.05 with |logLFC| > 0.3 threshold for color). *NAT10* is highlighted (bold) as a significantly upregulated ac4C writer (logLFC = 0.50, padj = 0.0023); the top three ranking hits by padj (OSGEPL1, THUMPD2, TARBP1) are labeled for reference. (C) *NAT10* expression (TPM) in liver samples from spaceflight (Space vs. Ground). Boxes show the interquartile range (IQR); horizontal lines, medians; individual data points superimposed. Group differences were assessed using the Wilcoxon rank-sum test.

Next, we performed correlation analyses between *NAT10* and functionally related genes. We selected a panel of 42 genes based on their reported or putative functional associations with NAT10-related pathways, including translation initiation and elongation factors, mRNA metabolism enzymes (e.g., decapping, deadenylation, and degradation), stress response genes (e.g., ATF4 and *DDIT3*), ribosome quality control components, and known *NAT10* targets or interactors identified in the literature (Supplementary Table S1). This list was curated to represent a functionally diverse but mechanistically relevant set of NAT10-associated processes. *NAT10* expression showed consistent positive correlations with *XRN1*, *EIF4G1*, and *DDIT3* expression in FLT samples (Fig. 2, Supplementary Fig. 2). *DDX17* was independently identified as a top-ranked NAT10-correlated gene in the genome-wide correlation analysis described below (Fig. 3), rather than as part of this curated 42-gene panel. These results suggest that *NAT10* is associated with RNA metabolic processes under spaceflight-induced stress conditions.

**Figure 2.**
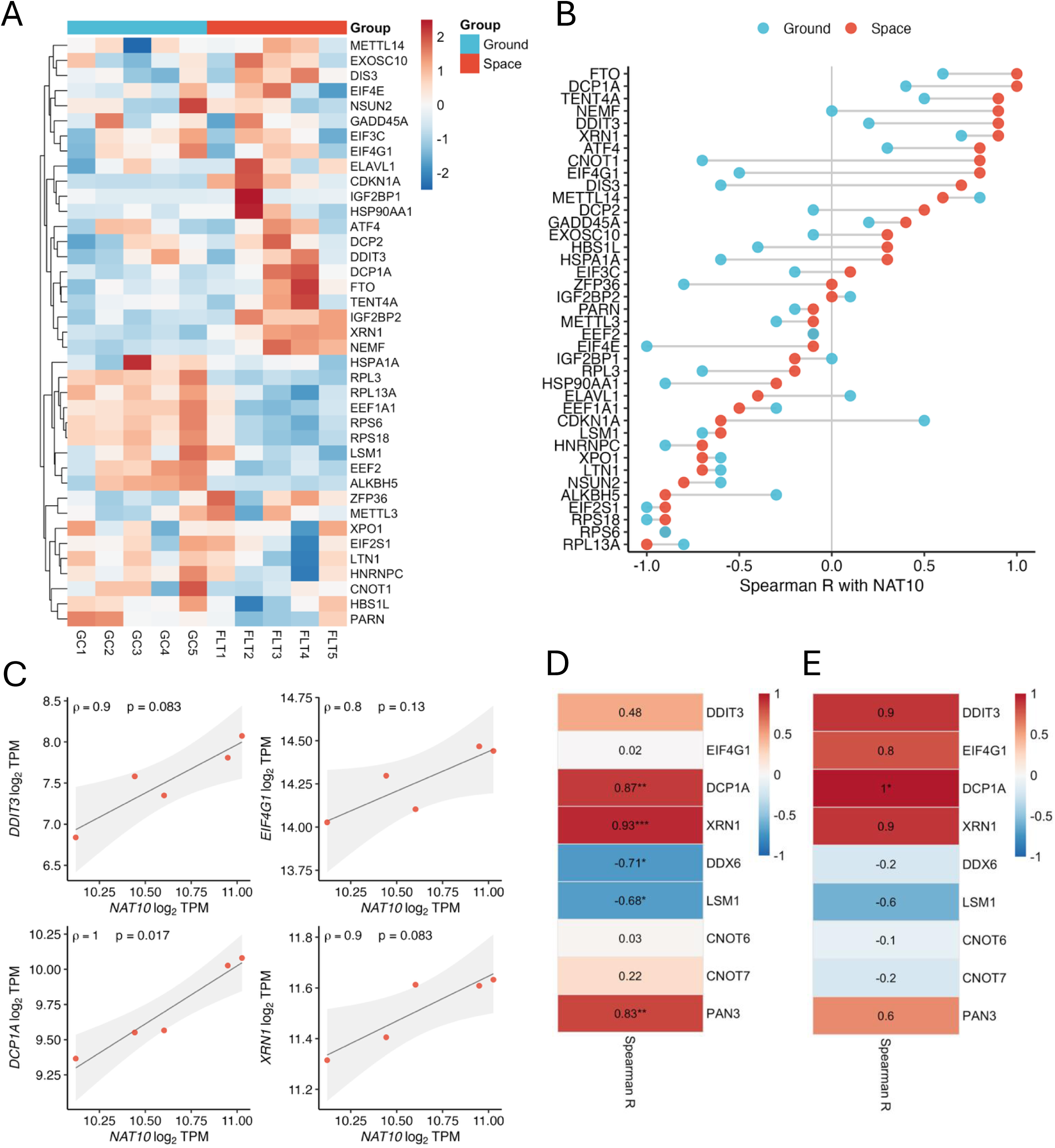
*NAT10* co-expression network implicates P-body and mRNA decay machinery. **(A)** Z-score heat map of genes coexpressed with *NAT10* in the liver (top-ranked by Spearman correlation with *NAT10*; Space + Ground samples, n = 10). Columns are ordered as Ground (n = 5), then Space (n = 5); rows are clustered by hierarchical clustering. Color scale: z-score of log₂ TPM. **(B)** Lollipop chart of Spearman correlation coefficients (ρ) between *NAT10* and co-expressed genes in Space (filled circles, red) and Ground (filled circles, teal) samples. The genes are ranked by Space correlation. Connecting lines span the Space–Ground difference. **(C)** Scatter plots of *NAT10* log₂ TPM versus log₂ TPM of four key RNA stability factors (*DDIT3*, *EIF4G1*, *DCP1A*, *XRN1*) in Space liver samples (n = 5). Linear regression with a 95% confidence band (gray) is shown. Spearman ρ and p-value are annotated. **(D)** Spearman correlation heat map between *NAT10* and P-body/mRNA decay factors (*DDIT3*, *EIF4G1*, *DCP1A*, *XRN1*, *DDX6*, *LSM1*, *CNOT6*, *CNOT7*, *PAN3*) in the liver samples (Space + Ground; n = 10). Color scale: Spearman ρ. **(E)** Same as (D) using Space liver samples only (n = 5).

**Figure 3.**
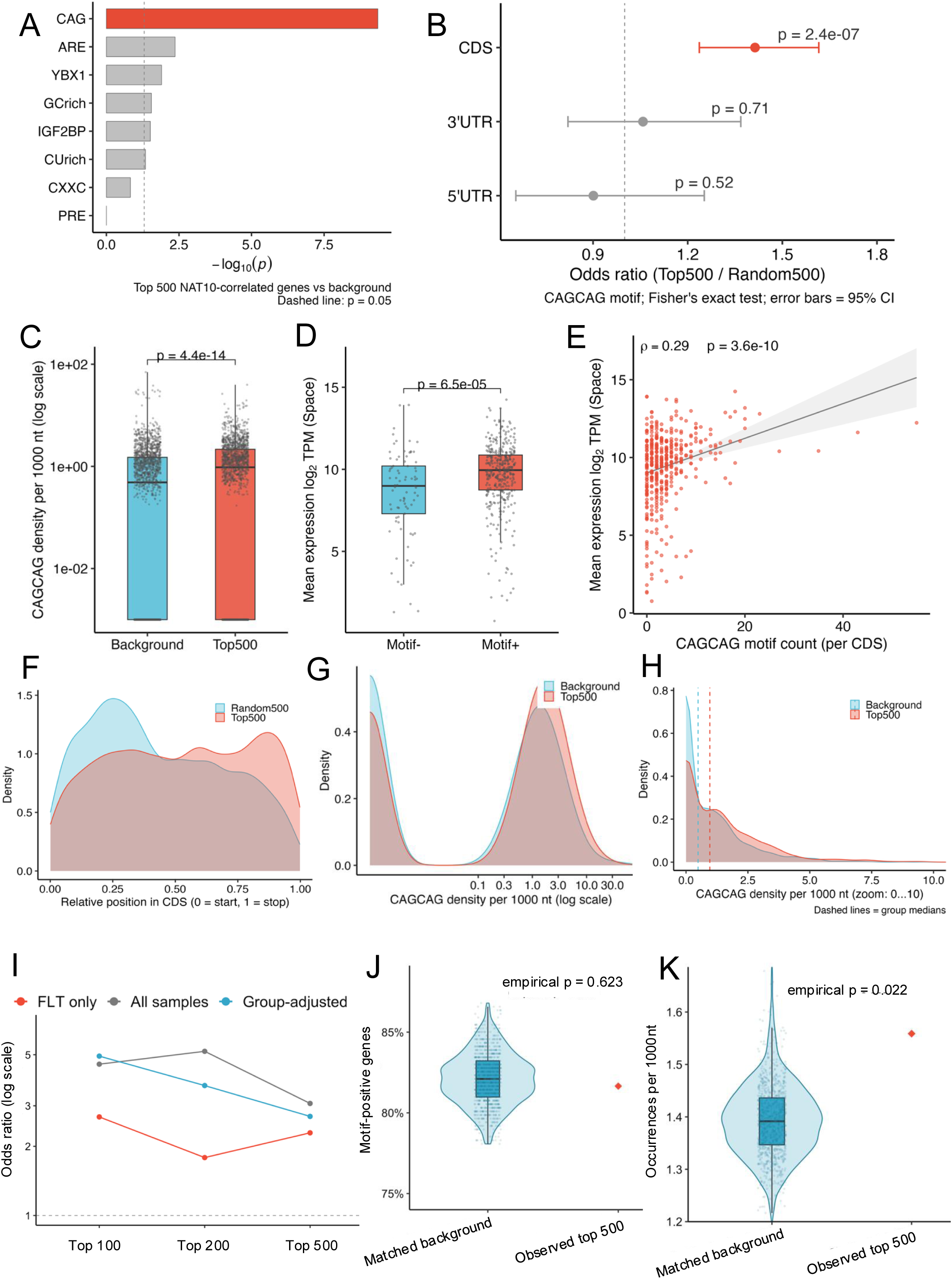
CAGCAG hexamer motif density is enriched in *NAT10*-correlated mRNAs. **(A)** Bar plot of −log₁₀(p) values from Fisher’s exact tests for enrichment of specific motifs (CAG, ARE, YBX1, GC-rich, IGF2BP, CU-rich, CXXC, and PRE) in the top-500 *NAT10*-correlated genes versus 500 random background genes. The dashed line indicates p = 0.05. **(B)** Forest plot of the odds ratios for CAGCAG motif enrichment stratified by genomic region (3′UTR, 5′UTR, CDS) in top-500 *NAT10*-correlated genes. Points, odds ratio; horizontal lines, 95% confidence intervals; dashed line at odds ratio = 1. **(C)** Boxplot of CAGCAG motif density (occurrences per 1,000 nt, log₁₀ scale) for the top-500 *NAT10*-correlated genes (red) versus 500 random background genes (teal). Statistical differences were evaluated using the Wilcoxon rank-sum test. **(D)** Boxplot of mean spaceflight expression (log₂ TPM) comparing CAGCAG motif-positive (Motif+) with motif-negative (Motif−) genes. Statistical differences were evaluated using the Wilcoxon rank-sum test. **(E)** Scatter plot of CAGCAG motif count per CDS versus mean log₂ TPM expression in Space liver samples. Each point represents one gene; linear regression with a 95% confidence band is shown. **(F)** Kernel density plot of the relative CAGCAG motif position along the CDS (0 = start codon, 1 = stop codon) in the top-500 (red) versus random-500 (teal) genes. **(G)** Density distribution of CAGCAG motif density (occurrences per 1,000 nt) plotted on a log₁₀ x-axis, comparing the top-500 *NAT10*-correlated genes (red) versus random-500 background genes (teal). **(H)** Same as (G) with x-axis limited to 0–10 to highlight low-density distributions. **(I)** Odds ratios for enrichment of CAGCAG motif-positive genes among the top 100, 200, and 500 *NAT10*-correlated genes. Correlations were calculated using spaceflight samples only (FLT only), all liver samples (FLT and GC), or all liver samples after subtracting the mean expression within each experimental group (group-adjusted). The dashed line indicates an odds ratio of 1. **(J)** Fraction of genes containing at least one CAGCAG motif in the observed top-500 *NAT10*-correlated gene set compared with 1,000 background gene sets matched for CDS length, GC content, and mean expression in FLT liver. The violin and boxplot show the distribution of the matched backgrounds; the red diamond represents the single observed summary statistic calculated for the Top-500 gene set. Empirical p = 0.623. **(K)** Mean CAGCAG motif density in the observed top-500 *NAT10*-correlated gene set compared with 1,000 background gene sets matched for CDS length, GC content, and mean expression in FLT liver. Motif density was calculated as the number of CAGCAG occurrences per 1,000 nt of CDS. The violin and boxplot show the distribution of the matched backgrounds; the red diamond indicates the observed top-500 value. Empirical p = 0.022.

### CAG repeat motif-containing transcripts are enriched among *NAT10*-correlated genes

To identify potential downstream targets of *NAT10*, we scanned for sequence motifs enriched among NAT10-correlated genes in the liver under spaceflight conditions. CAGCAG motif density was enriched within coding sequences of the top 500 NAT10-correlated genes in FLT liver samples (Fig. 3A, B, J, K). Initial comparisons with randomly sampled coding sequences highlighted CAG repeats relative to other candidate RNA motifs, including AU-rich elements (AREs) and PREs (Fig. 3A).

Unmatched comparisons showed higher CAGCAG motif frequency and density among NAT10-correlated genes than among random background genes (Fig. 3C, F–H). After matching background genes for CDS length, GC content, and expression, binary CAGCAG motif presence was not enriched, whereas motif density remained modestly but significantly elevated (empirical p = 0.022; Fig. 3J, K). Exploratory analyses also showed higher expression among CAGCAG motif-positive genes and a positive association between motif count and expression (Fig. 3D, E). Thus, motif density, rather than binary motif presence, was the more robust sequence-level association.

### *NAT10*-correlated genes are functionally linked to RNA metabolic processes

Gene Ontology (GO) analysis demonstrated marked enrichment of *NAT10*-correlated genes in RNA metabolic functional categories, encompassing splicing, catabolism, and processing (Fig. 4A–C). These functions were further enriched when examining CAG repeat motif-containing subsets of *NAT10* targets, consistent with an association between CAG-rich transcripts and RNA-homeostasis functions during spaceflight (Fig. 4D). Consistently, mean expression of CAG-repeat motif-positive genes associated with *NAT10* was positively correlated with individual RNA metabolic process genes, reaching nominal significance for *DCP1A* (Spearman ρ = 1.0, exact p = 0.017) with a similar trend for *XRN1* and *DDIT3* (ρ = 0.9, exact p = 0.083 each) at this sample size (n = 5; Fig. 4E). GO analysis supported enrichment of RNA metabolism-related functions (Fig. 4F), whereas the exploratory GSEA did not reach statistical significance for the pathways shown in Fig. S3A–D.

**Figure 4.**
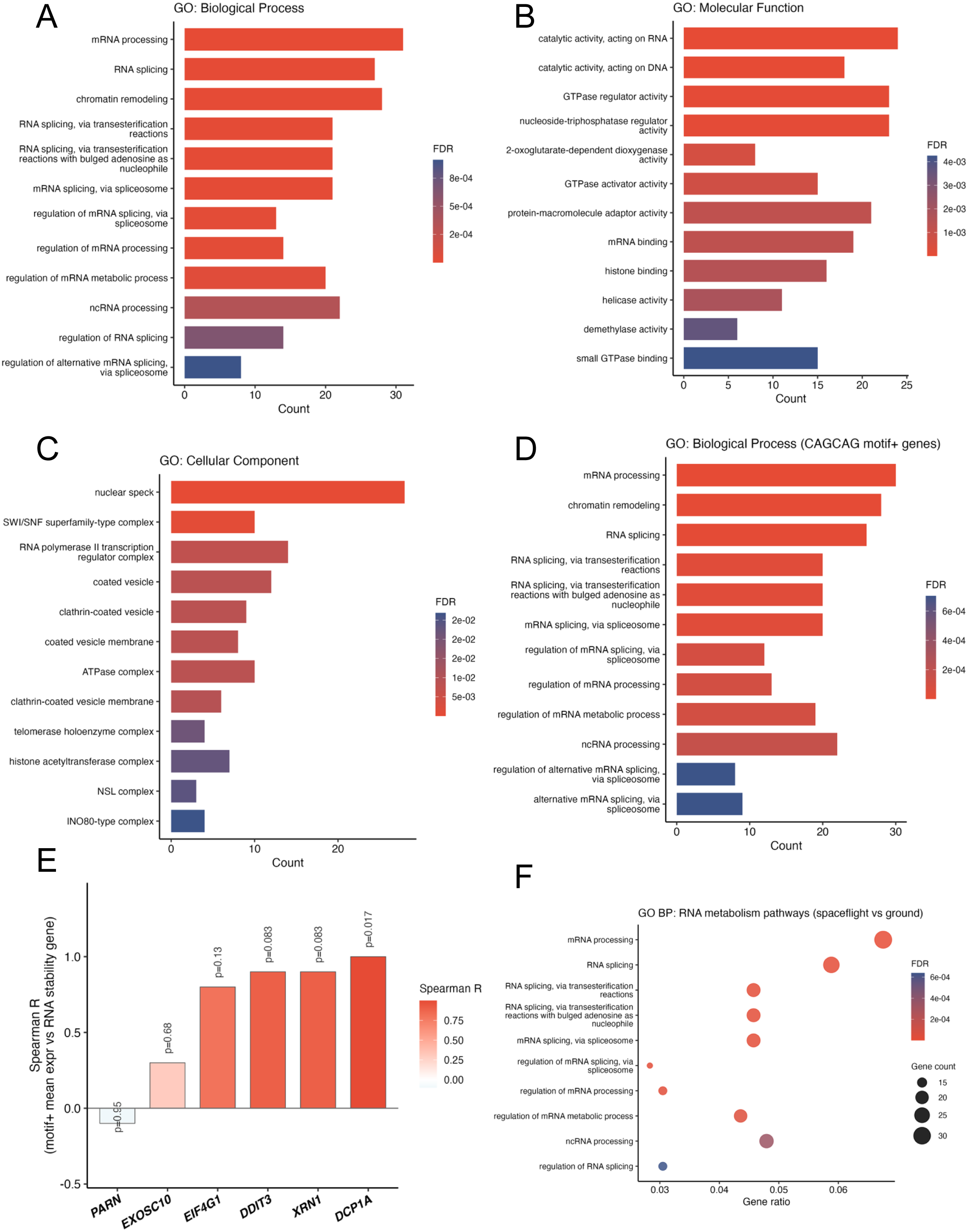
*NAT10*-correlated and CAGCAG motif-positive genes are enriched for RNA metabolism functions. **(A–C)** Gene Ontology (GO) enrichment barplots for Biological Process (A), Molecular Function (B), and Cellular Component (C) terms in the top-500 *NAT10*-correlated genes. The top 12 terms ranked by adjusted p-value are shown. Bar color: FDR (gradient from red to navy). Enrichment was performed using clusterProfiler. **(D)** GO Biological Process enrichment barplot for CAGCAG motif-positive genes (Motif+, n = top 500 *NAT10*-correlated genes). Same display as (A). **(E)** Spearman correlation barplot between the mean expression of CAGCAG motif-positive genes and individual RNA stability factor genes (*DCP1A*, *XRN1*, *DDIT3*, *EIF4G1*, *EXOSC10*, *PARN*) among Space liver samples (n = 5). Bar fill: Spearman ρ (blue–white–red gradient). p-values are annotated above each bar. **(F)** GO Biological Process dot plot for RNA metabolism-related pathways (filtered from the top-500 GO BP results). Dot size: gene count; dot color: FDR.

### NAT10-motif associations are partially shared across tissues

To assess the tissue specificity of NAT10-associated motifs, we compared their expression and correlation patterns across the liver, kidney, retina, heart, muscle, and spleen. Although *NAT10* expression was highest in the liver, the positive correlations between *NAT10* and RNA metabolism-related genes were observed across several tissues (Fig.5B,C, Supplementary Fig. 4). Moreover, we also observed positive correlations between *NAT10* and genes containing CAG repeat, CXXC, and IGF2BP motifs in five tissues for which CAGCAG motif enrichment data were available (Liver, Kidney, Heart, Spleen, and Retina; Fig. 5D). These findings suggest that the NAT10-motif axis may represent a recurring but tissue-dependent association during the spaceflight response.

**Figure 5.**
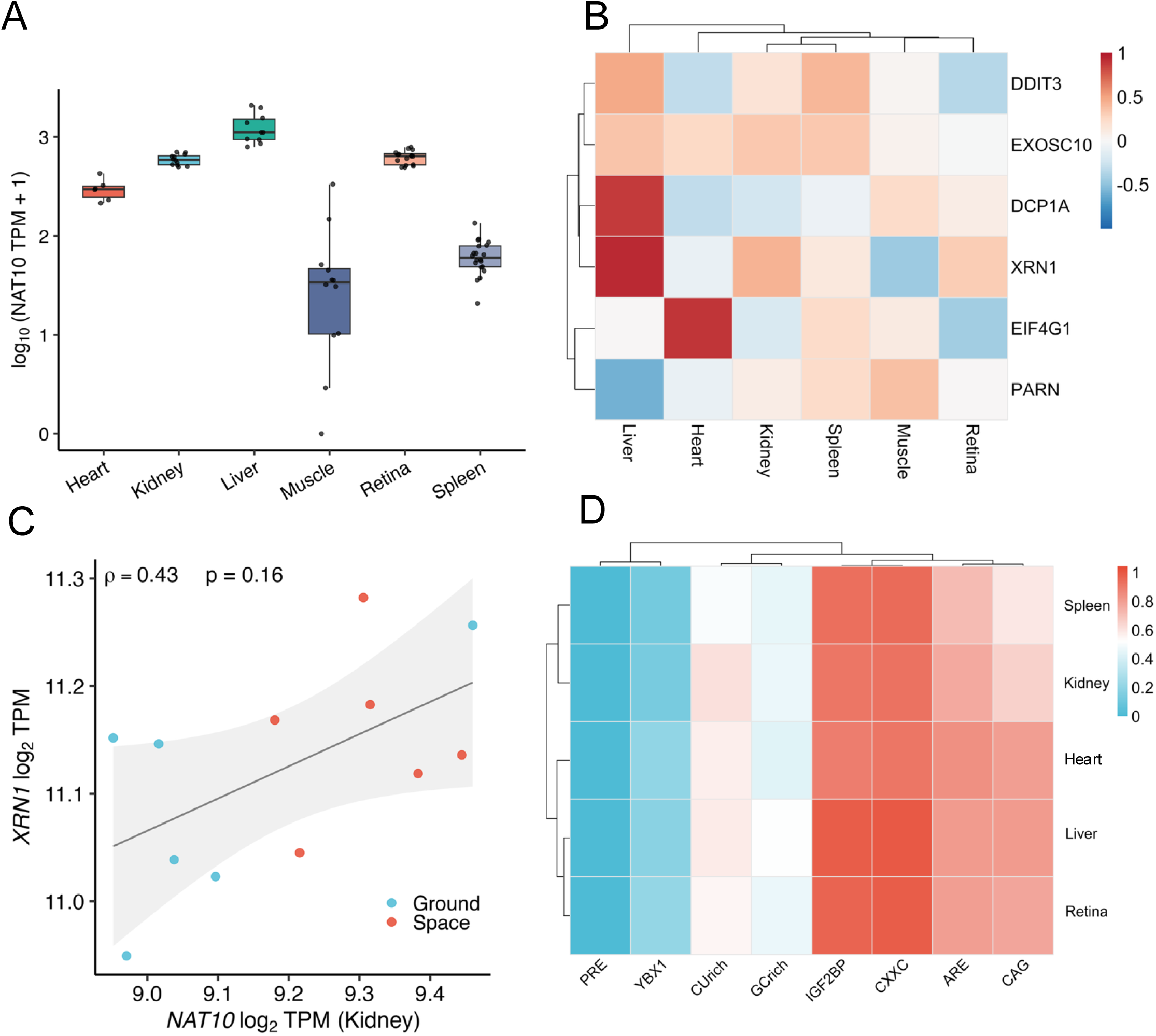
*NAT10*–mRNA decay co-expression is partially shared across multiple tissues. **(A)** Boxplot of *NAT10* expression [log₁₀(TPM + 1)] among six tissues (heart, kidney, liver, muscle, retina, spleen) from NASA GeneLab (all groups combined). Each data point represents one sample. Colors indicate tissue. **(B)** Spearman correlation heat map between *NAT10* and six RNA stability genes (*DCP1A*, *XRN1*, *DDIT3*, *EIF4G1*, *EXOSC10*, *PARN*) among six tissues. Correlation was computed per tissue using all available samples (Space + Ground). Color scale: Spearman ρ. **(C)** Scatter plot of *NAT10* versus *XRN1* log₂(TPM + 1) in Kidney samples (all groups; Space in red, Ground in teal). Linear regression with 95% confidence band is shown. Spearman ρ and p-value are annotated. **(D)** CAGCAG motif enrichment heat map (Top-200 correlated genes per tissue, fraction of genes containing ≥1 CAGCAG motif) across five tissues for which CAGCAG motif data were available (Liver, Kidney, Heart, Spleen, and Retina). Muscle was omitted due to insufficient sample availability for the multi-tissue motif analysis pipeline. Rows and columns are hierarchically clustered.

### Spaceflight-associated *NAT10* expression correlates with hepatic metabolic programs

Given that *NAT10*-correlated genes were enriched in RNA metabolic processes, including mRNA decay and processing, we hypothesized that these changes may have downstream consequences for hepatic metabolic homeostasis. To evaluate this, we applied GSEA to the liver transcriptome, contrasting FLT with GC conditions. GSEA indicated significant suppression of fatty acid β-oxidation (HALLMARK_FATTY_ACID_METABOLISM: NES = −1.98, FDR = 0.014; REACTOME_MITOCHONDRIAL_FATTY_ACID_BETA_OXIDATION: NES = −2.02, FDR = 0.023) and oxidative phosphorylation (HALLMARK_OXIDATIVE_PHOSPHORYLATION: NES = −2.46, FDR = 0.014) in FLT liver samples (Fig. 6A). Conversely, cholesterol biosynthesis pathways were significantly enriched in FLT samples (REACTOME_CHOLESTEROL_BIOSYNTHESIS: NES = +2.16, FDR = 0.023; REACTOME_REGULATION_OF_CHOLESTEROL_BIOSYNTHESIS_BY_SREBP_SR EBF: NES = +2.10, FDR = 0.023; HALLMARK_CHOLESTEROL_HOMEOSTASIS: NES = +1.63, FDR = 0.020). These findings are consistent with previously reported spaceflight-associated fatty liver phenotypes and indicate a shift in hepatic lipid metabolism under spaceflight conditions.

**Figure 6.**
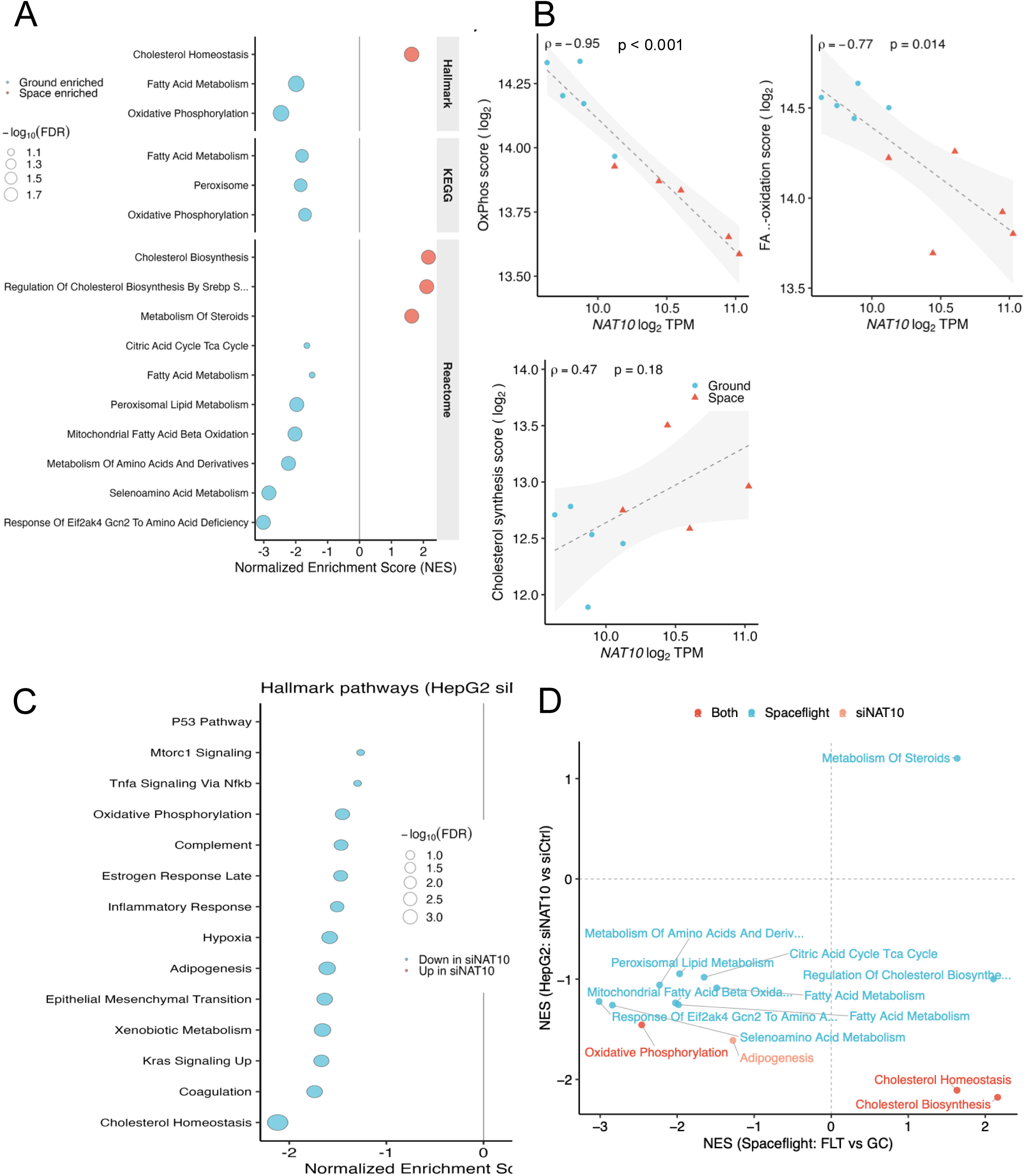
Spaceflight-associated *NAT10* expression correlates with hepatic metabolic programs, and *NAT10* depletion selectively opposes the cholesterol program. **(A)** Gene Set Enrichment Analysis (GSEA) dot plot for metabolic pathways enriched in liver (Space vs. Ground; NASA GeneLab). Only pathways with an FDR < 0.1 matching metabolic terms (fatty acid oxidation, oxidative phosphorylation, cholesterol, TCA cycle, etc.) are displayed, faceted by gene set database (Hallmark, KEGG, Reactome). Dot size: −log₁₀(FDR); fill: enrichment direction. **(B)** Scatter plots of *NAT10* logL TPM versus mean metabolic gene scores, oxidative phosphorylation (OxPhos), fatty acid β-oxidation (FA β-ox), and cholesterol biosynthesis (Chol), in Liver samples (n = 10). Scores were calculated as the mean logL TPM of representative gene sets. Space (red triangles) and Ground (teal circles) are shown separately. Spearman ρ and p-value (n = 10, FLT + GC) are annotated. Within FLT samples alone (n = 5, exact two-tailed permutation p-values): OxPhos ρ = −1.00, p = 0.017; FA β-ox ρ = −0.20, p = 0.783; Chol ρ = +0.30, p = 0.683. **(C)** GSEA dot plot for hallmark gene sets in HepG2 cells with *NAT10* knockdown (siNAT10 vs. siCtrl RNA-seq). Pathways with FDR < 0.25 are shown. Dot size: −log₁₀(FDR); fill: enrichment direction. *NAT10* knockdown efficiency was confirmed by RNA-seq (log₂FC = −1.65, FDR < 0.001). **(D)** NES mirror scatter plot comparing normalized enrichment scores (NES) for metabolic pathways between the spaceflight dataset (x-axis: FLT vs. GC) and the *NAT10* knockdown dataset (y-axis: siNAT10 vs. siCtrl). Points are colored by significance: red (Both datasets, FDR < 0.1 spaceflight and FDR < 0.25 siNAT10), teal (Spaceflight only), salmon (siNAT10 only), gray (not significant in either). Selected pathways are labeled.

Within the liver transcriptome (n = 10; FLT = 5, GC = 5), *NAT10* levels exhibited pronounced inverse correlations with key oxidative phosphorylation genes, including Atp5a1 (Spearman’s ρ = −0.988, p < 0.0001), Cox6a1 (ρ = −0.927, p = 0.0001), Atp5b (ρ = −0.927, p = 0.0001), and Sdha (ρ = −0.879, p = 0.002), as well as fatty acid β-oxidation genes Cpt2 (ρ = −0.770, p = 0.014) and Cpt1a (ρ = −0.709, p = 0.028; Fig. 6B). In contrast, *NAT10* expression positively correlated with the cholesterol biosynthesis gene Dhcr7 (ρ = +0.661, p = 0.044). These correlations, computed across all liver samples (n = 10), are consistent with the GSEA findings. To assess whether they reflected within-spaceflight covariation or between-group separation, Spearman correlations were re-computed in FLT samples only (n = 5). The inverse correlation with oxidative phosphorylation score persisted within FLT samples alone (ρ = −1.00, exact p = 0.017, two-tailed permutation test), although the small sample size (n = 5) warrants cautious interpretation, whereas the FA β-oxidation score correlation was substantially attenuated and not significant within FLT alone (ρ = −0.20, exact p = 0.783), suggesting that the FA β-oxidation association primarily reflects between-group (FLT vs. GC) differences rather than within-spaceflight coregulation.

To assess whether *NAT10* was functionally associated with these metabolic changes, we performed RNA-seq analysis of HepG2 hepatocellular carcinoma cells subjected to *NAT10* siRNA knockdown (siNAT10, n = 4) or nontargeting control siRNA (siCtrl, n = 4). We confirmed the *NAT10* mRNA knockdown using differential expression analysis (log₂FC = −1.65, FDR < 0.001). GSEA of Hallmark gene sets indicated that cholesterol homeostasis was among the most significantly suppressed pathways following *NAT10* depletion (NES = −2.12, FDR = 3.2×10⁻L), accompanied by suppression of adipogenesis (NES = −1.61, FDR = 0.010), xenobiotic metabolism (NES = −1.66, FDR = 0.010), and oxidative phosphorylation (NES = −1.45, FDR = 0.037; Fig. 6C). A NES mirror plot comparing the spaceflight and siNAT10 GSEA profiles showed that cholesterol biosynthesis (Reactome NES: spaceflight = +2.16, siNAT10 = −2.18) and cholesterol homeostasis (Hallmark NES: spaceflight = +1.63, siNAT10 = −2.11) were enriched in the spaceflight transcriptome and concordantly suppressed following *NAT10* depletion in HepG2 cells (Fig. 6D). Although transcriptome profiling alone cannot establish causality, these concordant patterns in a hepatocellular carcinoma cell line are consistent with a role for *NAT10* in hepatic lipid metabolism and raise the possibility that *NAT10* upregulation contributes to the metabolic reprogramming observed under spaceflight conditions.

### *NAT10* associates with and stabilizes CAG repeat-containing mRNAs

To dissect the enzymatic domain of *NAT10* required for mRNA stabilization, we expanded the actinomycin D-based mRNA stability assay to five conditions: control siRNA (siCtrl), *NAT10* knockdown (siNAT10), siNAT10 co-transfected with wild-type *NAT10* (siNAT10L+LNAT10-WT), siNAT10 with the acetyltransferase-dead G641E mutant (siNAT10L+LNAT10-G641E), and siNAT10 with the K290A helicase-domain mutant (siNAT10L+LNAT10-K290A). siRNA targeted the 3′UTR of endogenous *NAT10*, rendering CDS-only rescue constructs siRNA-resistant. *DDX17* mRNA, whose coding sequence contains three CAGCAG hexamers, was assayed by actinomycin D (ActD) chase over an extended 0/4/8 h time course. siNAT10 numerically decreased *DDX17* mRNA remaining at 8 h relative to siCtrl (17.1L±L1.3% vs. 44.4L±L13.1%), although this difference did not reach statistical significance by Dunnett’s test (pL=L0.128; one-way analysis of variance (ANOVA), FL=L7.15, pL=L0.0055; Fig.L7A, B): a trend was present, but statistical significance was not reached for this comparison. Co-transfection with wild-type *NAT10* significantly restored *DDX17* mRNA stability relative to siNAT10 (66.8L±L10.8% remaining; pL=L0.006 by Dunnett’s test), and the acetyltransferase-dead NAT10-G641E mutant produced a comparable, statistically significant rescue (63.8L±L7.8% remaining; pL=L0.009 vs. siNAT10), consistent with acetyltransferase activity not being strictly required under these conditions; whether the G641E rescue reflects residual catalytic activity or differences in mutant abundance, stability, or localization relative to wild-type *NAT10* remains unresolved. In contrast, the NAT10-K290A helicase-domain mutant failed to restore mRNA stability (24.5L±L1.3% remaining; pL=L0.919 vs. siNAT10), remaining statistically indistinguishable from siNAT10 alone and consistent with a contribution of helicase-domain integrity to NAT10-dependent *DDX17* mRNA stabilization. To assess the contribution of one selected site, the assay was repeated with a *DDX17* construct in which one of the three CAGCAG hexamers was deleted (DDX17-ΔCAG); the other two sites remained intact. DDX17-ΔCAG mRNA showed no significant differences in stability across any of the five conditions (67.8%, 61.3%, 70.8%, 50.6%, and 60.0% remaining at 8 h for siCtrl, siNAT10, +NAT10-WT, +NAT10-G641E, and +NAT10-K290A, respectively; one-way ANOVA, FL=L0.26, pL=L0.898; Fig.L7A, B). Despite the presence of two additional CAGCAG sites, no significant differences among *NAT10* conditions were detected for the DDX17-ΔCAG reporter, consistent with a role for this CAGCAG-containing region in NAT10-dependent stabilization, although the difference in rescue magnitude between the WT and ΔCAG reporters was not directly tested (see Limitations).

To determine whether this helicase domain-dependent pattern extends to additional NAT10-correlated transcripts, we conducted the same five-condition actinomycin D assay for *DCP1A*, a NAT10-correlated gene whose coding sequence contains four CAGCAG hexamers. At 8 h after actinomycin D treatment, *DCP1A* transcript abundance trended lower in siNAT10 cells (40.7L±L7.1% remaining) compared with siCtrl (64.7L±L1.4% remaining), although this difference did not reach statistical significance (one-way ANOVA, FL=L3.29, pL=L0.058; Dunnett’s test, siCtrl vs. siNAT10, pL=L0.750; Fig.L7A). Re-expression of wild-type *NAT10* numerically restored *DCP1A* mRNA stability (96.6L±L16.5% remaining; pL=L0.150 vs. siNAT10), and NAT10-G641E showed a similar trend toward rescue (107.6L±L34.3% remaining; pL=L0.075 vs. siNAT10), whereas NAT10-K290A did not (35.2L±L9.9% remaining; pL=L0.998 vs. siNAT10). This pattern is directionally consistent with the *DDX17* results—wild-type *NAT10* and the acetyltransferase-dead G641E mutant trending toward rescue while the K290A helicase-domain mutant does not—although the *DCP1A* comparisons did not reach statistical significance at this sample size (nL=L3): a similar trend was present, but statistical significance was not reached at this sample size, particularly among the rescue conditions. The DCP1A-ΔCAG construct, in which one of four CAGCAG hexamers was deleted, showed no significant differences in mRNA stability across any of the five conditions (57.8%, 50.7%, 62.4%, 55.2%, and 58.4% remaining at 8 h for siCtrl, siNAT10, +NAT10-WT, +NAT10-G641E, and +NAT10-K290A, respectively; one-way ANOVA, FL=L0.14, pL=L0.962; Fig.L7B), consistent with the absence of significant condition-dependent differences observed for the DDX17-ΔCAG reporter. Taken together, the *DDX17* and *DCP1A* single-site deletion results support a functional contribution of the selected CAGCAG-containing regions to NAT10-associated mRNA stabilization: neither ΔCAG reporter showed significant differences among the tested *NAT10* conditions despite retaining additional CAGCAG sites elsewhere in the coding sequence. The *DDX17* findings reached statistical significance for the NAT10-WT and NAT10-G641E rescue conditions, whereas the corresponding *DCP1A* trends, though directionally consistent, will require additional biological replicates to establish statistical significance. The G641E rescue reached statistical significance for *DDX17* but only a non-significant trend for *DCP1A*; taken together, these observations are consistent with acetyltransferase activity not being strictly required for NAT10-dependent mRNA stabilization under these conditions; whether it reflects a fully ac4C-independent mechanism, residual catalytic activity, or differences in mutant expression, stability, or localization has not been directly tested.

To determine whether deletion of selected CAGCAG sites alters NAT10-associated RNA recovery, we conducted RNA immunoprecipitation (RIP) assays with cells expressing NAT10-FLAG, followed by reverse-transcription quantitative PCR (RT-qPCR) for *DDX17* and *DCP1A*. As initial evidence, FLAG-tagged NAT10-WT preferentially enriched wild-type *DDX17* mRNA over DDX17-ΔCAG (Fig. 7C), with a directionally consistent pattern observed for DCP1A-WT over DCP1A-ΔCAG. These results support preferential *NAT10* association with reporters containing intact CAGCAG regions, as deletion of selected sites markedly reduced NAT10-associated RNA recovery.

**Figure 7.**
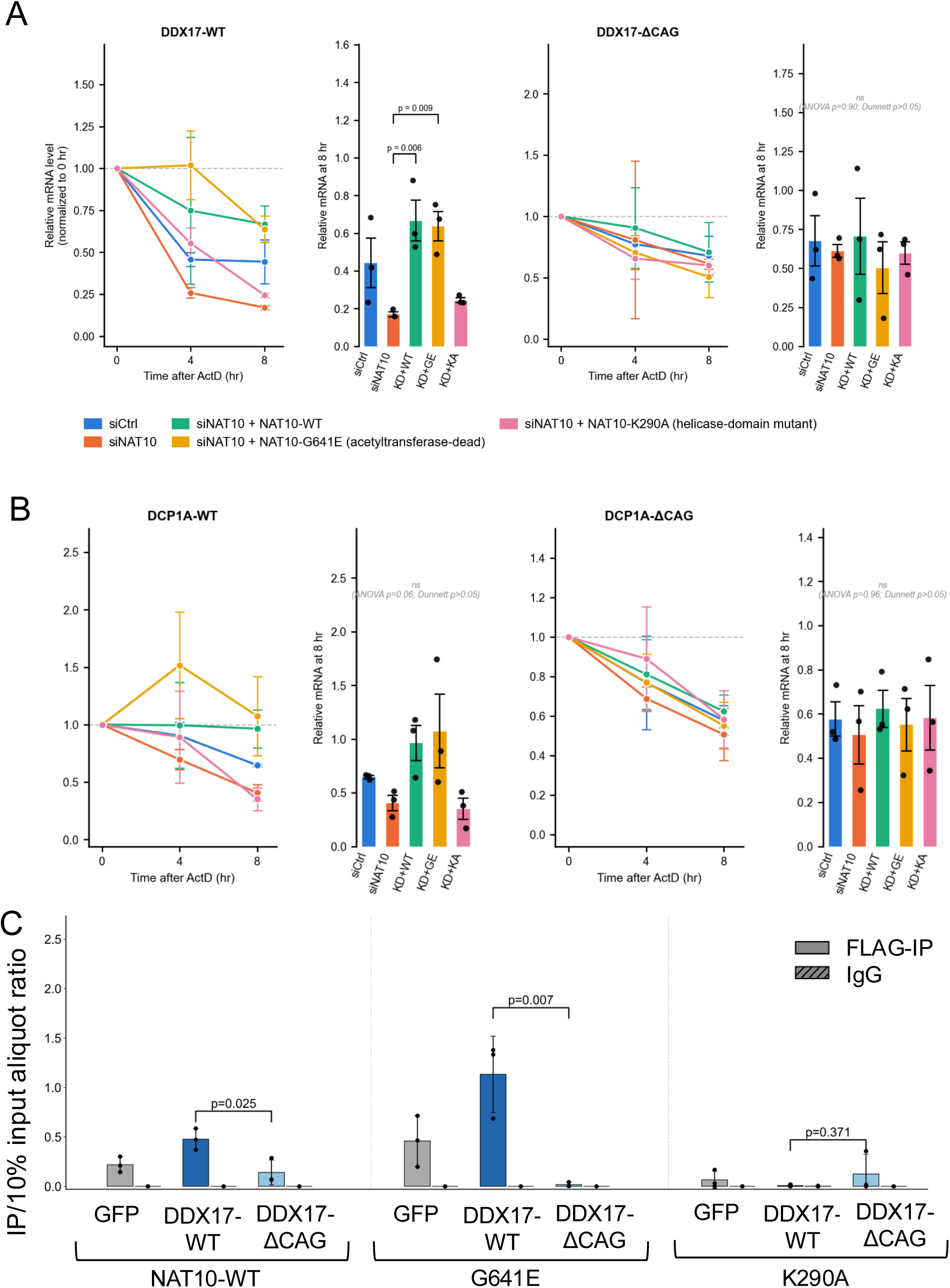
*NAT10* regulates mRNA stability through the CAGCAG motif and sequence-dependent RNA association. (A, B) Actinomycin D (ActD) mRNA decay assay in HepG2 cells. mRNA levels of DDX17-WT, DDX17-ΔCAG, DCP1A-WT, and DCP1A-ΔCAG reporter constructs (CDS only; DDX17-ΔCAG deletes one of three CAGCAG sites, whereas DCP1A-ΔCAG deletes one of four) were quantified by RT-qPCR at 0, 4, and 8Lh after ActD treatment under five conditions: siCtrl, siNAT10, siNAT10L+LNAT10-WT (wild-type rescue), siNAT10L+LNAT10-G641E (acetyltransferase-dead rescue), and siNAT10L+LNAT10-K290A (helicase-domain-mutant rescue). siRNA targeted the endogenous *NAT10* 3′UTR; rescue constructs encode CDS only and are siRNA-resistant. RT-qPCR values were normalized to mature 18S rRNA and subsequently expressed relative to the corresponding 0Lh time point. Left panels: mRNA decay curves; right panels: bar graphs at 8Lh. Data are meanL±LSEM (nL=L3). Statistical significance at 8Lh was assessed by one-way ANOVA followed by Dunnett’s multiple comparisons test against the siNAT10 (KD) group. DDX17-WT (one-way ANOVA FL=L7.15, pL=L0.0055): siCtrl vs. siNAT10, pL=L0.128 (NS); siNAT10+NAT10-WT vs. siNAT10, pL=L0.006; siNAT10+NAT10-G641E vs. siNAT10, pL=L0.009; siNAT10+NAT10-K290A vs. siNAT10, pL=L0.919 (NS). DDX17-ΔCAG: no significant differences across conditions (one-way ANOVA FL=L0.26, pL=L0.898; all Dunnett comparisons vs. siNAT10, pL>L0.96). DCP1A-WT: a directionally consistent but statistically non-significant trend was observed (one-way ANOVA FL=L3.29, pL=L0.058); siCtrl vs. siNAT10, pL=L0.750; siNAT10+NAT10-WT vs. siNAT10, pL=L0.150; siNAT10+NAT10-G641E vs. siNAT10, pL=L0.075; siNAT10+NAT10-K290A vs. siNAT10, pL=L0.998 (NS). DCP1A-ΔCAG: no significant differences across conditions (one-way ANOVA FL=L0.14, pL=L0.962; all Dunnett comparisons vs. siNAT10, pL>L0.87). **(C)** RNA immunoprecipitation (RIP)-qPCR analysis of *DDX17* mRNA binding by FLAG-tagged *NAT10* domain mutants. HepG2 cells were co-transfected with one of three FLAG-tagged *NAT10* variants—FLAG-NAT10-WT, FLAG-NAT10-G641E (acetyltransferase-dead), or FLAG-NAT10-K290A (helicase-domain mutant)—together with one of three reporter plasmids per independent well: pcDNA-GFP (Ctrl), pcDNA-DDX17-WT (WT), or pcDNA-DDX17-ΔCAG (ΔCAG). Each of the nine conditions (3 *NAT10* variants × 3 reporters) was performed in three independent wells (nL=L3). Anti-FLAG immunoprecipitation (FLAG-IP, solid bars) and paired IgG immunoprecipitation (hatched bars) were conducted on the same cell lysate. *DDX17* mRNA in the WT and ΔCAG conditions was quantified using reporter-specific primers, whereas the Ctrl (pcDNA-GFP) condition, which has no co-transfected reporter, was quantified using endogenous-transcript-targeting primers (Methods); the Ctrl bars therefore report *NAT10* association with the endogenous *DDX17* transcript and should not be read as an assay-background control for the reporter-specific primers. *DDX17* mRNA enrichment is expressed as IP/Input ratio; DDX17-ΔCAG deletes one of three CAGCAG sites, with two sites remaining. Bars represent meanL±LSEM. DCP1A-ΔCAG data are excluded from this panel because only one of four CAGCAG sites was deleted and three remained. Student’s t-test comparing FLAG-IP of WT vs. ΔCAG reporter within each *NAT10* variant: NAT10-WT, pL=L0.025; G641E, pL=L0.007; K290A, pL=L0.371 (NS). Comparisons across *NAT10* variants for DDX17-WT reporter FLAG-IP: NAT10-WT vs. G641E, pL=L0.047; NAT10-WT vs. K290A, pL=L0.002.

To examine whether the failure of NAT10-K290A to rescue mRNA stability reflects impaired RNA-binding capacity, we performed RIP assays using FLAG-tagged NAT10-WT, NAT10-G641E, and NAT10-K290A constructs in HepG2 cells. FLAG-NAT10-WT enriched *DDX17* WT reporter mRNA at an IP/Input ratio of 0.477L±L0.107 (Fig.L7C). FLAG-NAT10-G641E showed at least comparable enrichment (IP/InputL=L1.133L±L0.386; pL=L0.047 vs. NAT10-WT), consistent with its retained mRNA-binding capacity and its rescue activity in the actinomycin D assays (comparable to wild-type for *DDX17*; a non-significant trend for *DCP1A*). In contrast, FLAG-NAT10-K290A exhibited significantly reduced enrichment of *DDX17* mRNA (IP/InputL=L0.008L±L0.011; pL=L0.002 vs. NAT10-WT) (Fig.L7C). Directionally consistent results were observed for *DCP1A* mRNA, with NAT10-WT showing the highest enrichment (IP/InputL=L0.164L±L0.037), NAT10-G641E showing intermediate enrichment (0.055L±L0.017), and NAT10-K290A showing minimal enrichment. These data are consistent with the K290A substitution impairing NAT10-associated RNA recovery and may help explain its reduced rescue capacity. Western blot analysis of the anti-FLAG immunoprecipitates confirmed comparable recovery of NAT10-WT, NAT10-G641E, and NAT10-K290A protein across all co-transfected reporter conditions (Supplementary Fig. S7D), indicating that the reduced RNA recovery observed for NAT10-K290A is not attributable to differential immunoprecipitation efficiency.

To explore whether deletion of the selected *DDX17* CAGCAG site could alter predicted RNA structure and *NAT10* contact geometry, we performed RNA secondary structure prediction and molecular docking simulations. RNAfold analysis of the *DDX17* CAG repeat region predicted distinct secondary structures for the wild-type (WT) sequence and the corresponding single-site ΔCAG construct (Fig. S6A–G). Molecular docking of the *NAT10* cryo-EM structure (PDB: 9B0E) with the WT *DDX17* CAG motif RNA identified 744 contact atoms within 5 Å, compared with 477 contact atoms for the ΔCAG RNA, yielding a larger predicted interface in the WT model than in the single-site deletion model. These structural models support the possibility that the selected CAGCAG-containing region influences RNA conformation and *NAT10* contact geometry, although binding affinity and direct sequence recognition remain to be tested experimentally.

To extend these findings to the endogenous context, we performed actinomycin D stability assays and RIP targeting endogenous *DDX17* and *DCP1A* transcripts without co-transfection of exogenous mRNA constructs. siNAT10 treatment accelerated *DDX17* mRNA decay compared with siCtrl cells (36.3% vs. 54.2% remaining at 4 h; p = 0.010; Fig. S7A), consistent with the overexpression rescue data. *DCP1A* mRNA showed a directionally consistent trend toward decreased stability in siNAT10 cells (35.3% vs. 51.5% remaining at 4 h), although statistical significance was not reached (p = 0.145; Fig. S7B; n = 3). Furthermore, expressed NAT10-FLAG RIP using an anti-FLAG antibody confirmed association with *DDX17* and *DCP1A* mRNAs (Fig. S7C). Together, these endogenous experiments support the relevance of NAT10-associated mRNA stabilization and RNA association to endogenous *DDX17* and *DCP1A* transcripts.

To further evaluate the cross-dataset reproducibility of NAT10-dependent metabolic regulation, we re-analyzed a publicly available *NAT10* knockdown RNA-seq dataset (GSE210086)^31^ generated in MCF7 cells. Using DESeq2, all eight fatty acid metabolism genes reported by Dalhat et al.^31^—ELOVL6, SCD, ACSL1, ACSL3, ACSL4, ACADSB, ACADM, and ACAT1—were consistently downregulated upon siNAT10 treatment (all p < 0.05; Fig. S8B). Gene Set Enrichment Analysis (GSEA) additionally indicated significant downregulation of HALLMARK_PROTEIN_SECRETION (NES = −1.85, FDR = 0.002) and HALLMARK_UV_RESPONSE_DN (NES = −2.23, FDR < 0.001; Fig. S8A), consistent with impaired lipid secretion programs. These cross-dataset findings provide independent consistency with NAT10-dependent fatty acid metabolic regulation beyond HepG2 cells, while not establishing equivalence across cell types. Published mouse-liver studies provide complementary evidence relevant to the cholesterol-related axis (Wang et al., 2024; Yang et al., 2024).^33,34^

## Discussion

Our systematic screen of RNA modification enzyme expression across spaceflight-exposed tissues identified *NAT10* among a set of substantially upregulated RNA modification-related factors in the liver, distinguished by a statistically robust induction in the liver, although not the numerically largest fold-change among the tissues examined. Transcriptome-wide correlation analysis associated *NAT10* expression with a module of RNA metabolism genes enriched for CAG repeat motifs in their coding sequences. Functional experiments implicated *NAT10* helicase domain integrity in the stability of CAG repeat-containing mRNAs: *NAT10* knockdown destabilized target transcripts, stability was restored to a comparable extent by wild-type *NAT10* and the acetyltransferase-dead G641E mutant, and not restored by the K290A helicase-domain mutant. RNA immunoprecipitation further showed that NAT10-associated RNA recovery was sensitive to single-site CAGCAG deletion in the tested reporters. In spaceflight liver, *NAT10* expression was also associated with suppression of fatty acid beta-oxidation and induction of cholesterol biosynthesis.

*NAT10* depletion selectively opposed the cholesterol-related, but not all, components of this metabolic signature in HepG2 cells. Together, these data support a model in which *NAT10* contributes to CAG repeat-associated mRNA stabilization, identified here as a candidate posttranscriptional mechanism of potential relevance to hepatic adaptation, alongside an independently observed association with cholesterol-related metabolic programs in spaceflight liver.

The enrichment of CAG repeat density and CXXC motifs among *NAT10*-correlated transcripts is consistent with sequence context influencing posttranscriptional target selectivity, in agreement with reports implicating the CXXC motif as an ac4C-recognition element ^11^. Although CAG repeats are best known for their roles in neurodegenerative disorders through polyglutamine expansion ^28,29^, CAG repeat-rich sequences may contribute to transcript stability and nuclear export ^22–27^, raising the possibility that these motifs serve as positive regulatory elements during spaceflight stress. The convergent *DDX17* stability and RIP results support a functional contribution of the selected CAGCAG-containing region to NAT10-dependent regulation. Because two other *DDX17* CAGCAG sites remained and deletion may also alter local RNA structure, direct recognition and motif sufficiency remain to be established. By contrast, the CXXC motif was not directly characterized at the biochemical level in the present study. Whether CXXC sequences represent additional binding or modification sites for *NAT10* warrants further investigation.

The functional enrichment of *NAT10*-correlated genes in RNA splicing, decay, and catabolic pathways aligns with the recognized function of *NAT10* in maintaining posttranscriptional homeostasis and with reports that ac4C-modified transcripts exhibit enhanced stability and translation efficiency ^11^. The further enrichment of CAG repeat motif-positive subsets within these pathways suggests that motif content confers an additional layer of selectivity during NAT10-associated posttranscriptional regulation.

IGF2BP-binding motifs were also enriched among *NAT10*-correlated genes. IGF2BP2, a well-characterized m6A reader, has been shown to stabilize transcripts under stress ^30^, and its coenrichment may reflect a layered posttranscriptional regulatory response.

The positive correlation between *NAT10* and RNA metabolism genes, and the enrichment of CAG repeat density in *NAT10*-correlated transcripts, were observed across multiple spaceflight-exposed tissues, including kidney, retina, heart, muscle, and spleen, even in tissues where *NAT10* was not substantially upregulated. This suggests that NAT10-associated RNA regulatory patterns occur within a recurring, tissue-dependent stress-associated module, potentially independent of expression alone. The degree of correlation varied by tissue; however, the pronounced hepatic upregulation of *NAT10* relative to other tissues may reflect the central role of the liver in lipid, cholesterol, and glucose homeostasis. Spaceflight-associated metabolic reprogramming, including the suppression of fatty acid β-oxidation and induction of cholesterol biosynthesis, may impart disproportionate posttranscriptional regulatory demands on hepatocytes, potentially contributing to the prominent *NAT10* response observed in this tissue.

Principal component analysis (PCA) of RNA modification enzyme expression profiles indicated that tissue identity is the primary driver of transcriptional variance, with spaceflight-associated (FLT vs. GC) differences emerging as a secondary pattern within each tissue (Supplementary Fig. 5A, B), consistent with tissue-specific baseline expression of RNA modification enzymes dominating global variance, whereas spaceflight exerts a more selective transcriptional impact. Differential expression analysis further identified the selective upregulation of enzymes associated with m5C, m6A, and ac4C modifications, suggesting coordinated changes in RNA-modification enzyme expression under spaceflight conditions.

Beyond RNA metabolism, suppression of fatty acid beta-oxidation and oxidative phosphorylation, together with induction of cholesterol biosynthesis in spaceflight-exposed liver, is reminiscent of a NAFLD-like hepatic phenotype reported in spaceflight models. The inverse correlation between *NAT10* expression and oxidative phosphorylation score persisted within FLT samples alone (ρ = −1.00, exact p = 0.017; n = 5, two-tailed permutation test), although the small sample size warrants cautious interpretation. By contrast, the FA β-oxidation score correlation was not significant within FLT samples alone (ρ = −0.20, exact p = 0.783), suggesting it primarily reflects between-group separation; this association should therefore be interpreted with caution. These correlations indicate association, not causation. In HepG2 cells, *NAT10* depletion opposed the spaceflight-associated cholesterol program, consistent with reports linking *NAT10* to cholesterol accumulation and fatty acid metabolic reprogramming in cancer cells ^31,32^. By contrast, oxidative phosphorylation was reduced in both spaceflight liver and NAT10-depleted cells. Thus, the present data support a selective relationship between *NAT10* and cholesterol-related pathways, whereas the broader metabolic response likely reflects additional spaceflight-induced mechanisms. Direct perturbation of *NAT10* in vivo will be required to test its causal contribution to hepatic metabolic adaptation.

Observations relevant to NAT10-dependent hepatic metabolic regulation have been reported in independent model systems (Supplementary Table S3). Wang et al. demonstrated that *NAT10* promotes liver lipogenesis through ac4C modification of Srebf1 and Scap mRNAs in AML12 mouse hepatocytes and HepG2 cells, with *NAT10* depletion reducing cellular lipid accumulation,^33^ corroborating the cholesterol-related effects observed in the present study. Yang et al. further showed that hepatocyte-specific *NAT10* knockout in mice improved MASLD and MASH phenotypes in vivo, including attenuation of hepatic lipid accumulation and metabolic dysregulation,^34^ providing direct causal evidence consistent with the present associative spaceflight data. In the hepatocellular carcinoma context, *NAT10* depletion suppressed proliferation and invasion in both HepG2 and Huh7 cells,^35^ demonstrating that *NAT10* function is not restricted to a single hepatic cell line. Furthermore, acRIP-seq analysis identified NAT10-dependent ac4C modification of target transcripts in HCC cell lines including HCCLM3,^36^ consistent with posttranscriptional regulatory activity of *NAT10* across hepatocyte-related cell types. Collectively, these independent reports from multiple hepatic model systems support the biological relevance of observations made in HepG2 cells, while not establishing HepG2 as fully representative of normal hepatocytes, and underscore the need for future validation using primary hepatocytes or hepatocyte-derived organoid systems.

The pattern observed across rescue and RIP assays is consistent with a model in which *NAT10* helicase-domain integrity and RNA association contribute to CAG repeat-associated mRNA stabilization. Wild-type *NAT10* and the acetyltransferase-dead G641E mutant rescued target mRNA stability to a comparable extent, whereas the K290A mutant failed to rescue and showed reduced RNA recovery in RIP assays. Preferential recovery of DDX17-WT over the single-site deletion reporter further supports a functional contribution of the selected CAGCAG-containing region to *NAT10*–RNA association. Western blot analysis confirmed comparable immunoprecipitation of NAT10-WT, NAT10-G641E, and NAT10-K290A protein (Supplementary Fig. S7D), arguing against differential immunoprecipitation efficiency as the explanation for the reduced RNA recovery observed for K290A. Similarly, whether the maintained G641E rescue observed for *DDX17* reflects a residual catalytic contribution of G641E or differences in mutant abundance, stability, or localization relative to wild-type *NAT10* remains unresolved.

The comparable rescue by G641E suggests that acetyltransferase activity is not strictly required for the stabilization response under the conditions tested. Arango et al. demonstrated that *DDX17* mRNA stability is NAT10-dependent by BRIC-seq analysis, yet *DDX17* was not recovered among direct ac4C modification sites in the same study,^11^raising the possibility that the G641E-dependent rescue reflects an indirect or cofactor-mediated mechanism rather than direct ac4C modification of *DDX17*. The downstream effector(s) remain to be identified. Helicase domains in RNA-processing factors can facilitate RNA strand separation, remodeling of RNA secondary structure, or displacement of competing RNA-binding proteins; resolving which of these activities participates in CAG repeat-associated mRNA stabilization will require CLIP-seq and biochemical analyses of relevant cofactors. A recent independent study reported that *NAT10* physically interacts with *DDX17* protein and regulates its expression, promoting tubular epithelial cell senescence in a cisplatin-induced acute kidney injury model^39^. This prior report already establishes a direct *NAT10*–*DDX17* relationship; the principal advance of the present work therefore lies not in identifying that *NAT10* and *DDX17* are functionally linked, but in showing that G641E-mediated rescue was maintained while K290A-mediated rescue was not observed, a pattern consistent with a role for the specific CAGCAG-containing region of *DDX17* mRNA and for *NAT10* helicase-domain integrity.

In summary, spaceflight transcriptomic data identified *NAT10* as a liver-upregulated candidate regulator associated with a CAG repeat-enriched RNA-metabolism gene module; functional experiments in HepG2 cells independently validated NAT10-dependent, helicase domain-associated stabilization of CAG repeat-containing mRNA (*DDX17*, with a directionally consistent trend for *DCP1A*); and spaceflight-associated hepatic metabolic changes provide related biological context whose mechanistic link to CAG repeat-associated mRNA stabilization remains to be established. This work broadens knowledge of posttranscriptional regulation associated with spaceflight adaptation and identifies *NAT10* and its associated RNA motifs as candidate regulators for future studies of long-term spaceflight biology.

### Limitations of the study

Although the spaceflight component of the present study was based primarily on analyses of publicly available transcriptomic datasets, additional experiments are needed to fully clarify the functional roles of *NAT10* and motif-enriched transcripts in the spaceflight context. Direct detection of ac4C modifications in mRNA substrates, *NAT10* perturbation experiments, and biochemical mapping of CAG repeat- and CXXC-containing transcripts would strengthen the proposed regulatory model substantially. Future studies incorporating targeted ac4C profiling and RNA stability assays are warranted to determine whether *NAT10* directly acetylates specific RNA motifs or exerts its effects through broader stress-adaptive mechanisms. Furthermore, whereas the primary liver differential expression comparison (Fig. 1A, B; Supplementary Fig. 5B) was reanalyzed from raw sequencing reads using a full STAR–featureCounts–DESeq2 pipeline, the multi-tissue and gene coexpression analyses (Fig. 2–6; Supplementary Figs. 1, 4, 5A) continue to rely on publicly available pre-normalized TPM matrices, and associated statistical comparisons should be interpreted as screening-level rather than definitive count-based inference for those analyses.

The initial enrichment of CAG repeat motifs in *NAT10*-correlated genes exceeded that in randomly sampled coding sequences (Fig. 3C, F–H). Sensitivity analyses provided a more nuanced interpretation. Enrichment of CAG-positive genes persisted after removal of the FLT-GC group mean, arguing against group separation as the sole explanation. However, after matching background genes for CDS length, GC content, and expression, motif presence alone was not increased, whereas motif density remained modestly elevated. Accordingly, we interpret CAG density, rather than the binary presence of a CAGCAG motif, as the more robust sequence-level feature. Because the FLT liver cohort contained five samples, individual gene ranks remain exploratory and require validation in independent spaceflight cohorts. Functional support was obtained by deleting one of three *DDX17* CAGCAG sites and one of four *DCP1A* sites. These experiments demonstrate contributions from selected CAGCAG-containing regions but do not distinguish direct sequence recognition from effects of motif position or local RNA structure, nor do they establish motif sufficiency or the functional consequences of transcript-wide motif density. Additionally, the conclusion that CAGCAG deletion abolishes condition-dependent rescue rests on separate one-way ANOVAs for each reporter (significant for the WT reporter, not significant for the ΔCAG reporter) rather than on a formal statistical test of a reporter×condition interaction; the two ANOVA outcomes are therefore suggestive of, but do not statistically establish, a difference in the magnitude of the rescue effect between reporters. Site-by-site and combinatorial analyses, together with formal interaction testing, will be useful to define how multiple CAGCAG sites cooperate.

Among characterized RNA modification enzymes, no acetyltransferase other than *NAT10* has been shown to install N4-acetylcytidine (ac4C) onto RNA, although the existence and functional significance of ac4C on mRNA are controversial ^16,17^. Because the present study did not directly assess ac4C levels, we interpreted the NAT10-dependent effects on mRNA stability and transcriptomic correlations as evidence of NAT10-dependent posttranscriptional function, while acknowledging that the precise molecular mechanism, whether through direct mRNA ac4C modification or through non-acetyltransferase roles, remains to be established. This distinction is deliberately reflected in our framing, which emphasizes NAT10-dependent, helicase domain-dependent RNA stabilization, rather than direct claims of mRNA acetylation.

## Acknowledgments

We thank the laboratory members of the Department of Systems Biomedicine, Institute of Science Tokyo, for their support, and Michiko Shoji for their kind support. This research was supported by JSPS KAKENHI (Grant numbers: 22K07226 and 25K10441 to R.K.), and grants from Mochida Memorial Foundation for Medical and Pharmaceutical Research, the Ichiro Kanehara Foundation, Japan Philanthropic Foundation, SGH Foundation, and MSD Life Science Foundation to R.K. The authors also thank Akira Kurimoto, Akane Kurimoto, and Shintaro Kurimoto for their curiosity and companionship during the early exploration of the NASA GeneLab database, which sparked the initial inspiration for this work.

## Author contributions

Conceptualization and research design: R.K. Data collection and in vitro experiments: R.K. Bioinformatic analysis: R.K. Manuscript preparation (original draft): R.K. Critical reading and revision: T.C., T.M., Y.U. Resource provision (laboratory equipment and instruments): T.T., I.O. Conceptual advice on metabolic analysis: T.T. Research environment: I.O. Supervision and scientific guidance: T.T., H.A. All authors read and approved the final version for submission.

## Figure legends

**Supplementary Figure S1.**
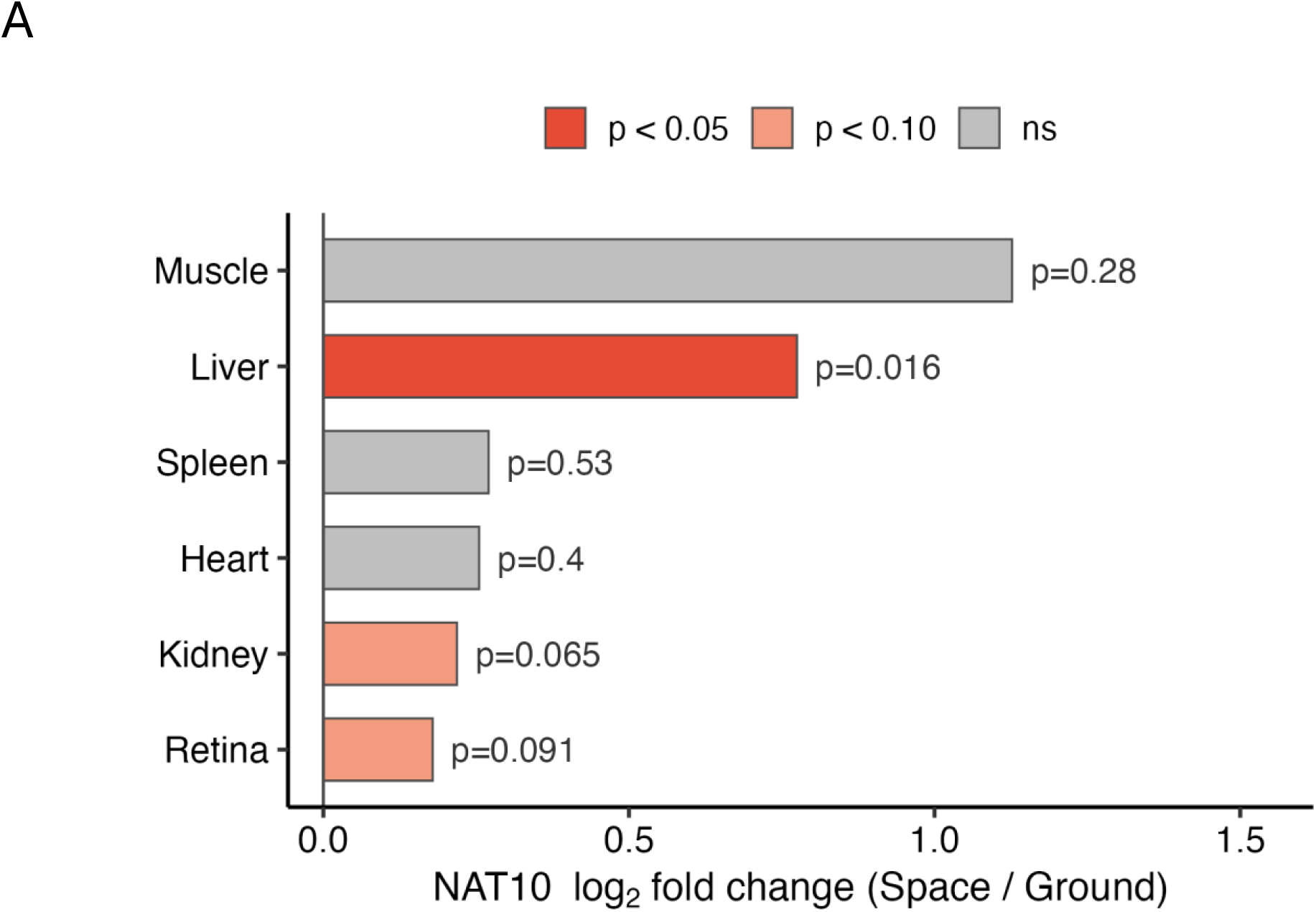
*NAT10* is upregulated in the spaceflight transcriptome across multiple tissues. Log₂ fold-change (Space vs. Ground) of *NAT10* expression across six tissues (heart, kidney, liver, muscle, retina, spleen) from NASA GeneLab datasets (GLDS-168, GLDS-420, GLDS-599, GLDS-419, GLDS-397, GLDS-102). Statistical significance between Space and Ground groups for each tissue was assessed by the Wilcoxon rank-sum test; FDR-adjusted p-values were calculated for reference.

**Supplementary Figure S2.**
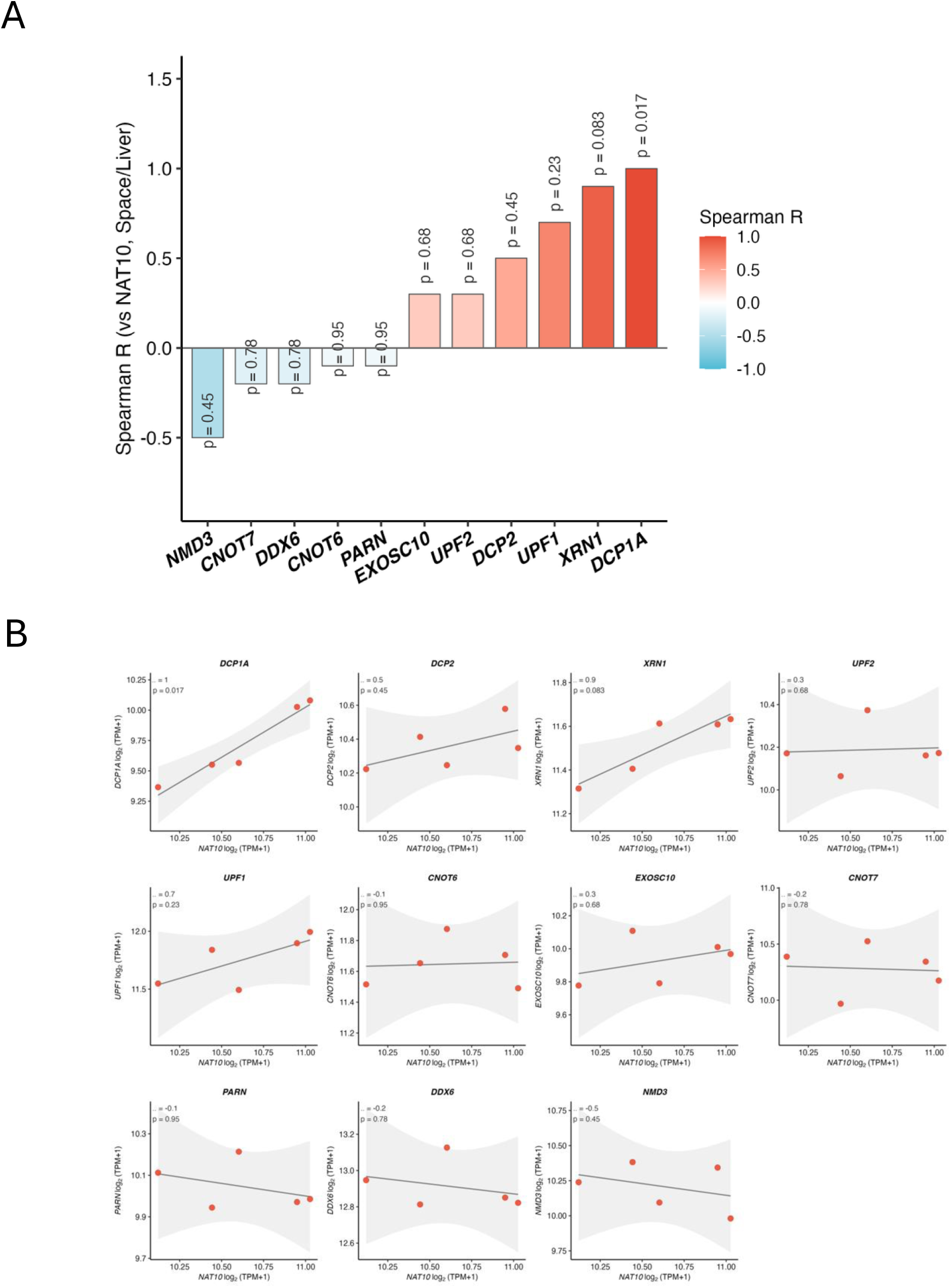
*NAT10* correlates with RNA stability factor expression in spaceflight liver. **(A)** Spearman correlation bar plot between *NAT10* and 11 RNA stability/decay factors (*DCP1A*, *DCP2*, *XRN1*, *EXOSC10*, *UPF1*, *UPF2*, *CNOT6*, *CNOT7*, *PARN*, *DDX6*, *NMD3*) in Space liver samples (FLT, n = 5). Bar fill: Spearman ρ. p-values are annotated above each bar. **(B)** Individual scatter plots of *NAT10* log₂(TPM + 1) versus each of the 11 RNA stability genes in Space liver samples (n = 5). Linear regression with 95% confidence band is shown. Spearman ρ and p-value are annotated in each panel.

**Supplementary Figure S3.**
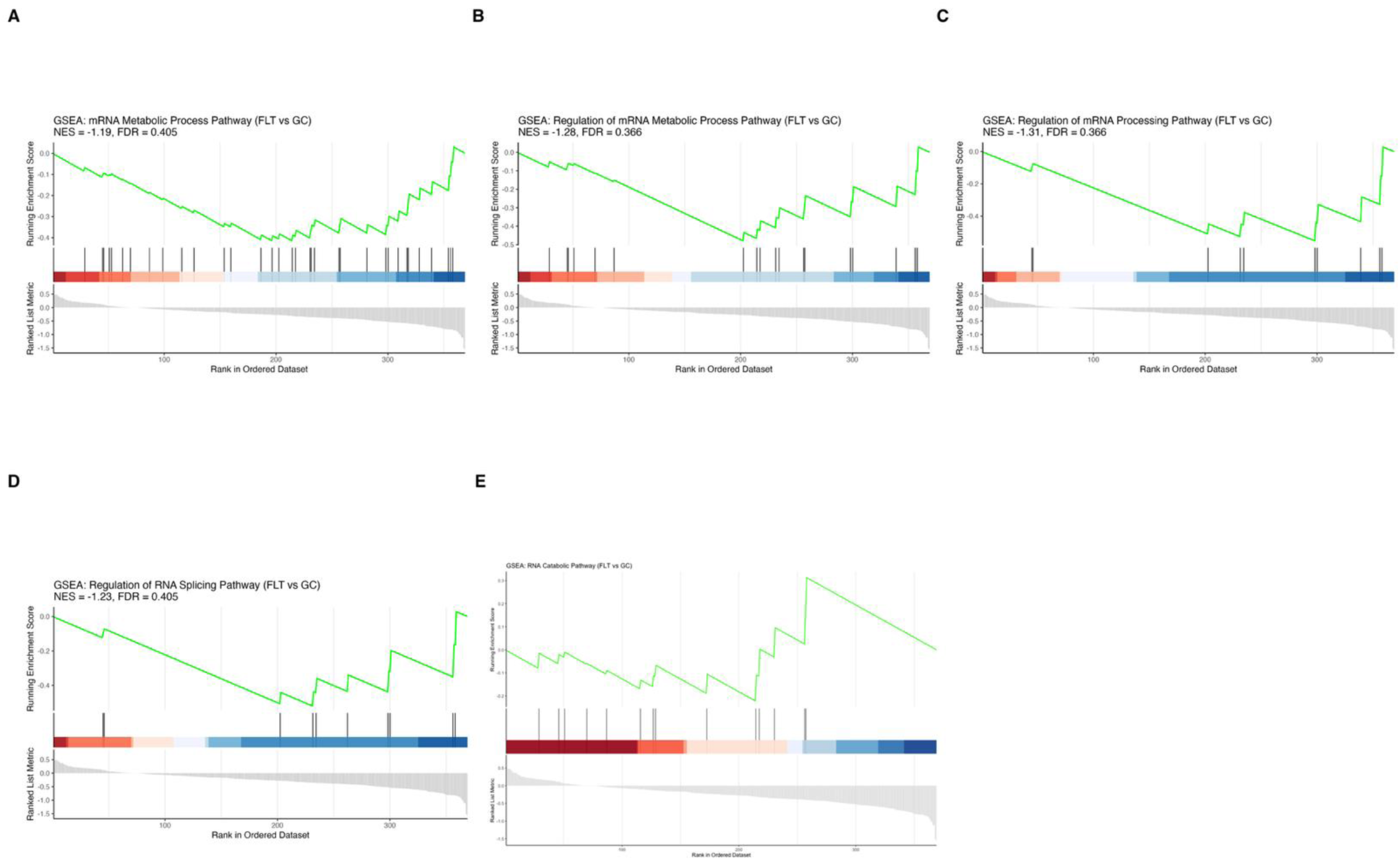
GSEA enrichment plots for RNA metabolic pathways in the spaceflight liver transcriptome. GSEA enrichment plots for five RNA metabolism gene sets: (A) mRNA Metabolic Process, (B) Regulation of mRNA Metabolic Process, (C) Regulation of mRNA Processing, (D) Regulation of RNA Splicing, and (E) RNA Catabolic Process. Gene sets are from Gene Ontology Biological Process (GO BP) via MSigDB (C5). NES and FDR for each gene set are shown in the title. The analysis was conducted using fgsea on gene expression fold-change (FLT vs. GC) rankings restricted to CAG repeat motif-positive genes (Space liver, NASA GeneLab). This exploratory analysis is presented for transparency; pathways in (A)–(D) did not reach statistical significance after FDR correction and should not be interpreted as evidence of significant enrichment.

**Supplementary Figure S4.**
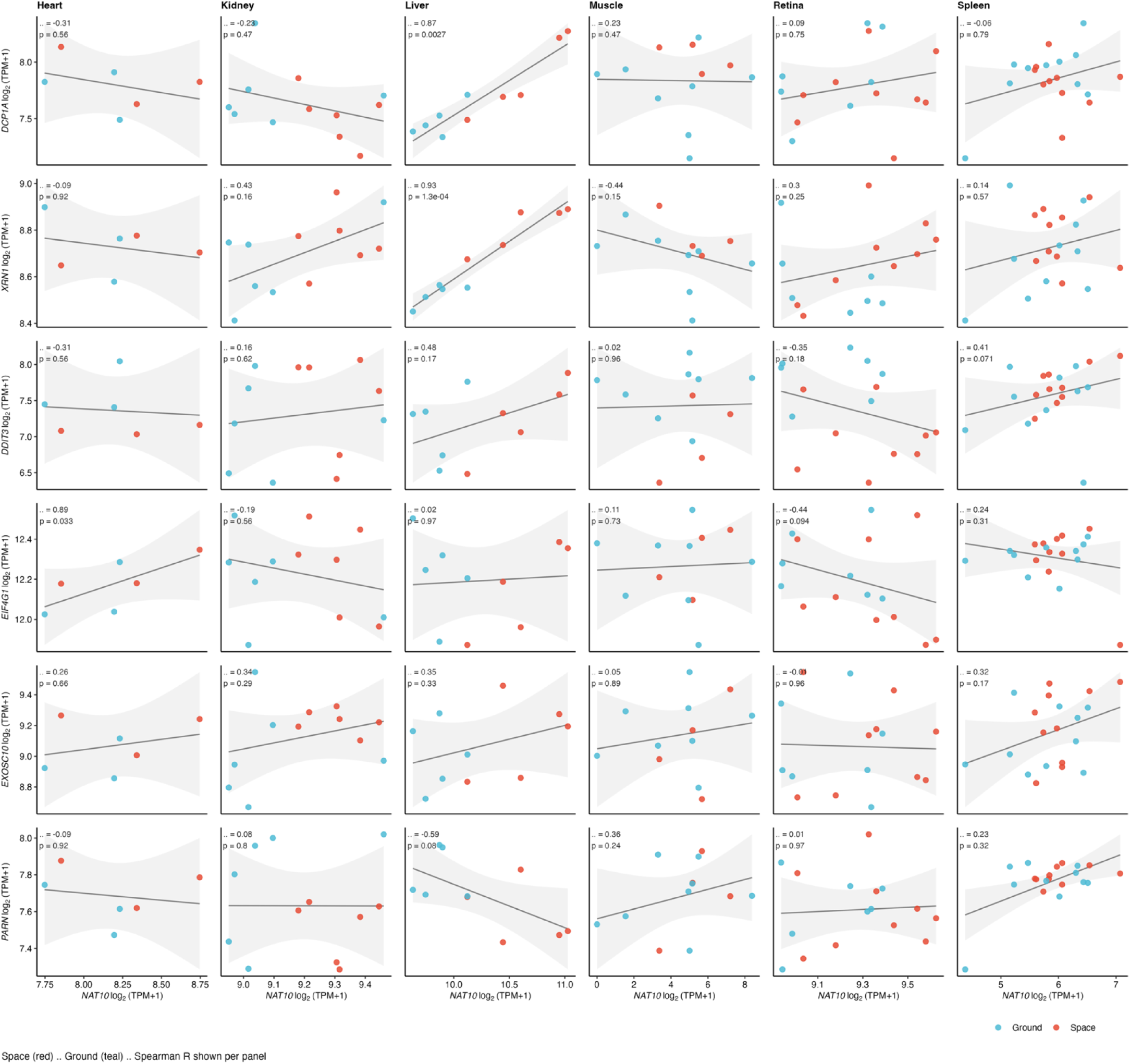
*NAT10*–mRNA decay gene correlations across tissues and spaceflight conditions. Scatter plots of *NAT10* versus six RNA stability gene (*DCP1A*, *XRN1*, *DDIT3*, *EIF4G1*, *EXOSC10*, *PARN*) log₂(TPM + 1) expression among six tissues (heart, kidney, liver, muscle, retina, spleen). Space samples are shown in red, and Ground samples are shown in teal. Rows correspond to genes; columns correspond to tissues (36 panels total). Linear regression with a 95% confidence band is shown per panel. Spearman ρ and p-value are annotated.

**Supplementary Figure S5.**
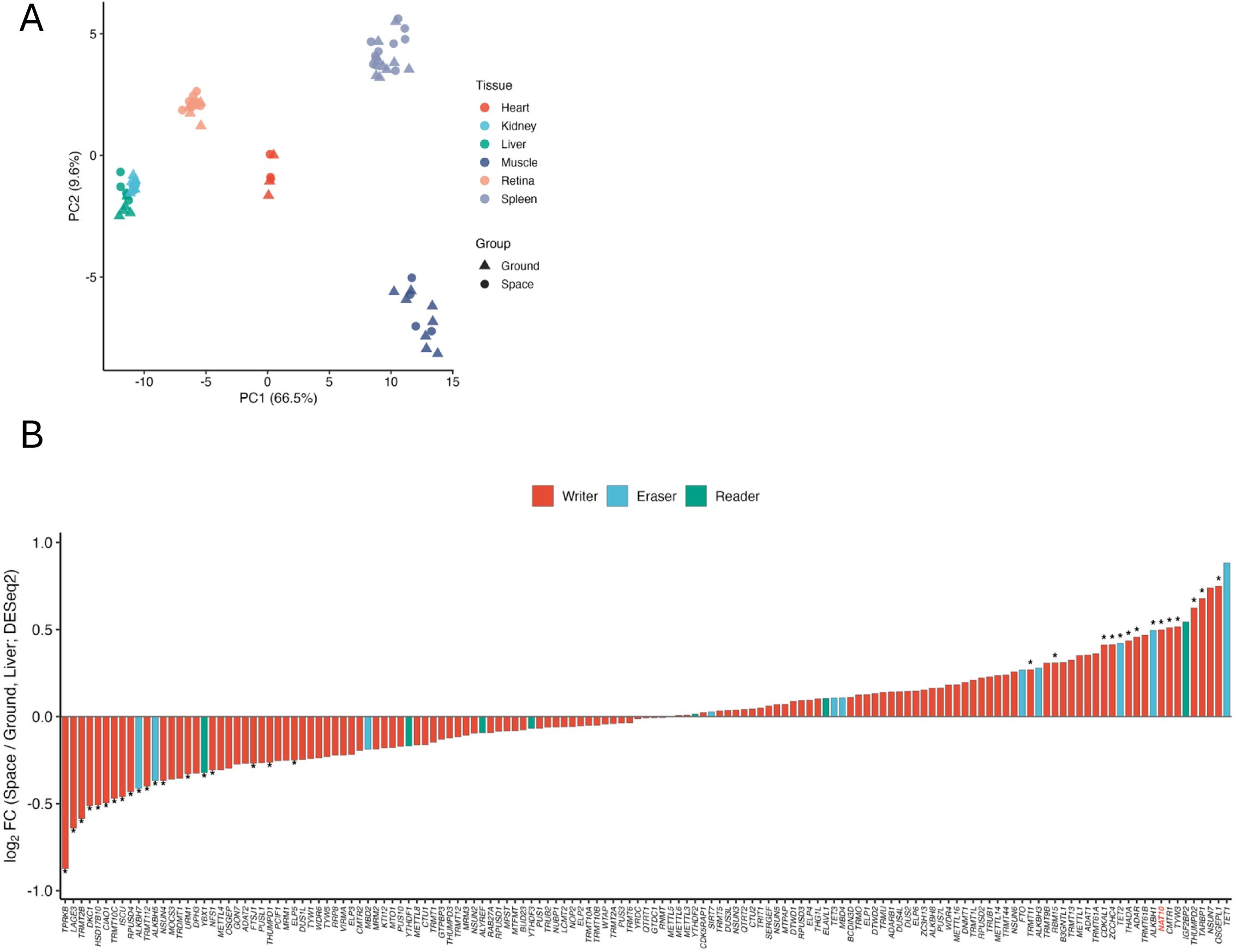
RNA modification enzyme expression profiles in spaceflight. **(A)** Principal component analysis (PCA) of RNA modification enzyme expression among all samples (n = 76; 6 tissues × Space and Ground). Gene features include writers, erasers, and readers spanning ac4C, m6A, m5C, m1A, m7G, pseudouridine (PseudoU), A-to-I editing, m6Am, and further tRNA/rRNA modification classes (e.g., t6A, m2G, 2’-O-methylation). The same curated panel of 152 RNA modification-related genes used in Fig. 1B and Supplementary Fig. 5B (n = 141 detected from 152 queried genes) was used for PCA visualization. Points are colored by tissue; shapes depict group (Space: circle, Ground: triangle). Variance explained by PC1 and PC2 is indicated on the axis labels. **(B)** LogL fold-change (Space vs. Ground, Liver; DESeq2) of the 152-gene RNA modification enzyme panel (143 genes detected), ranked in ascending order. Bar fill indicates enzyme type (Writer/Eraser/Reader); asterisks denote padj < 0.05. *NAT10* (ac4C, red label) showed a significant increase in spaceflight liver (logLFC = 0.50, padj = 0.0023) among a broader set of upregulated RNA modification enzymes, several of which (e.g., OSGEPL1/t6A, TET1/m5C, TARBP1/2’-O-methylation) exhibited numerically larger fold changes. RNA-seq data from spaceflight experiments were obtained from NASA GeneLab (GLDS-168, GLDS-420, GLDS-599, GLDS-419, GLDS-397, GLDS-102). Statistical tests and thresholds are as described for each panel. All box plots display median (center line), IQR (box), and 1.5× IQR (whiskers). Error bars represent SEM unless otherwise indicated.

**Supplementary Figure S6.**
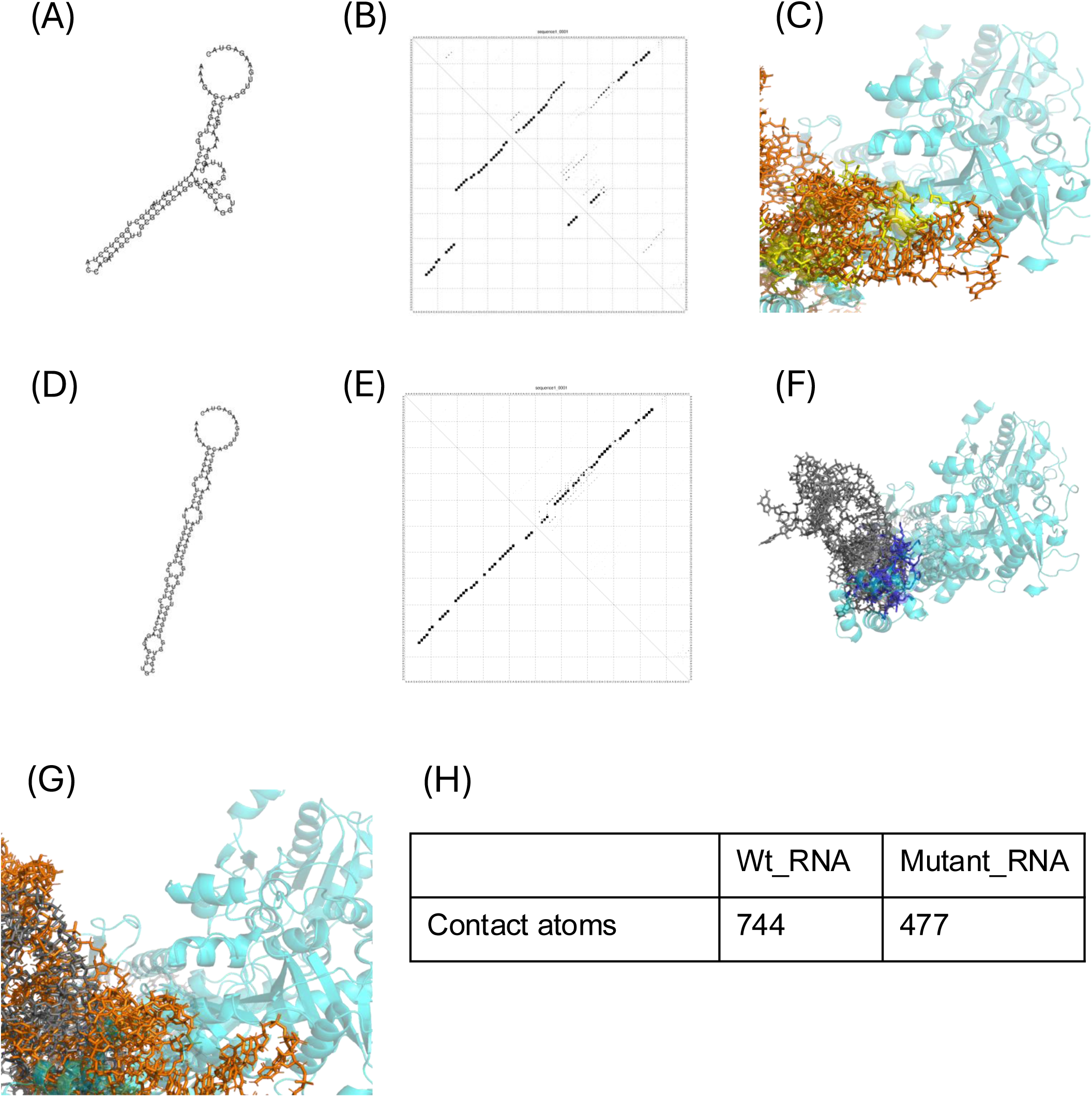
Structural modeling of *NAT10* interaction with a *DDX17* RNA region containing the selected CAGCAG site. (A–G) Predicted RNA secondary structures of the *DDX17* CAG repeat region: wild-type (WT, containing tandem CAG repeats; A–C) and single-site ΔCAG construct (one of three *DDX17* CAGCAG sites deleted; D–F). G: Merge images of A and D. Structures were predicted using RNAfold (ViennaRNA package); minimum free energy centroid structures are shown. Molecular docking simulation of *NAT10* protein (cyan ribbon; cryo-EM structure PDB: 9B0E) with the WT *DDX17* CAG motif RNA (orange sticks; left) and ΔCAG mutant RNA (gray sticks; right). Contact residues in *NAT10* within 5 Å of the RNA are indicated. The WT *DDX17* RNA model yielded 744 contact atoms compared with 477 for the single-site ΔCAG model, supporting the possibility that the selected region influences *NAT10*–RNA contact geometry; direct binding affinity remains to be tested experimentally. Visualization was conducted using PyMOL.

**Supplementary Figure S7.**
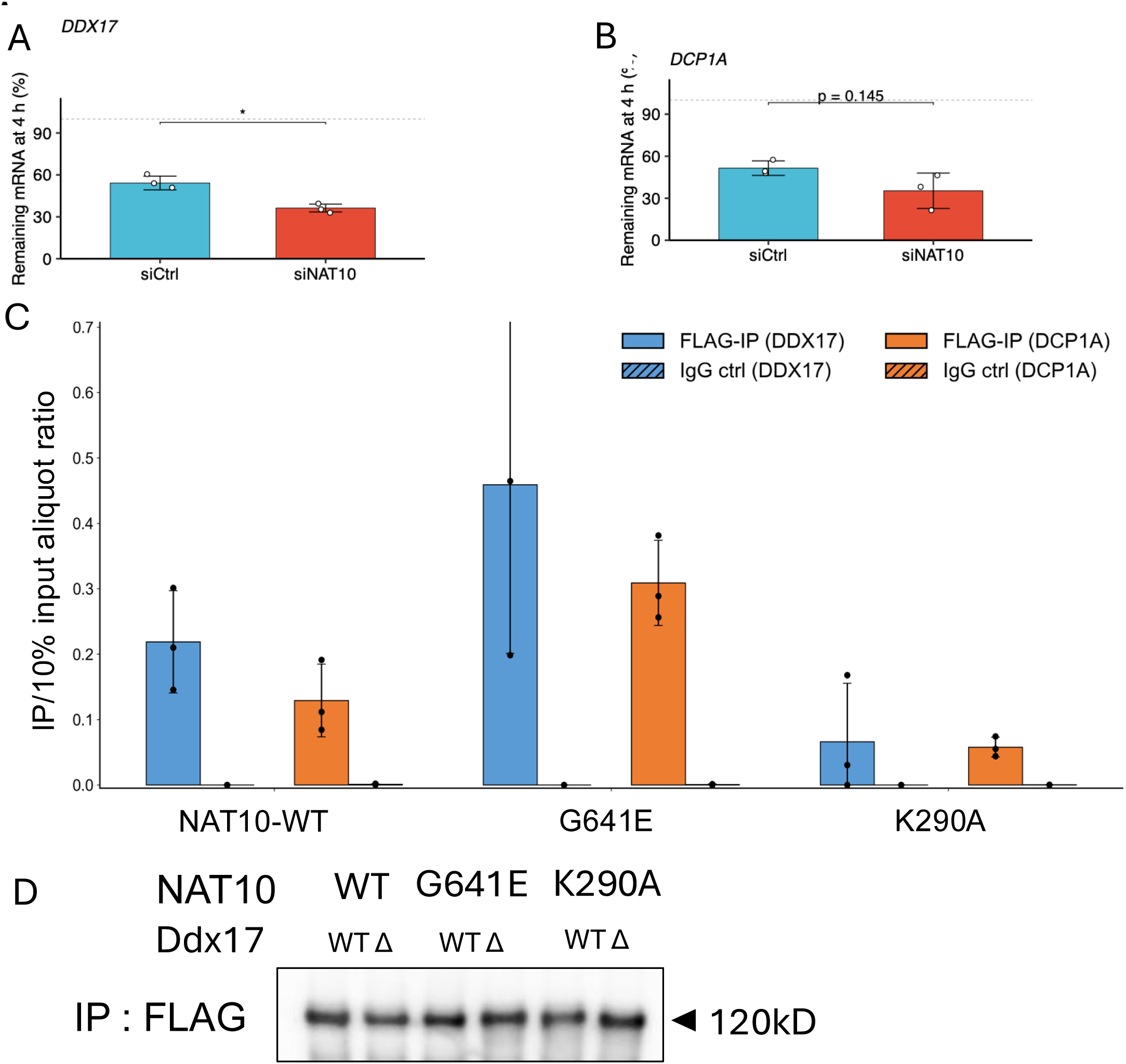
*NAT10* promotes endogenous mRNA stability and associates with target transcripts. (A–B) Actinomycin D (ActD) mRNA stability assay evaluating endogenous *NAT10* function. HepG2 cells were transfected with control siRNA (siCtrl) or siRNA targeting *NAT10* (siNAT10) 48 h prior to actinomycin D treatment (5 μg/mL). mRNA levels of *DDX17* and *DCP1A* were quantified by RT-qPCR at 0 and 4 h. RT-qPCR values were normalized to mature 18S rRNA and expressed as the percentage of remaining mRNA at 4 h relative to the corresponding 0 h time point (mean ± SD; n = 3). (A) *DDX17*: siCtrl 54.2 ± 4.9%, siNAT10 36.3 ± 2.8% (*p = 0.010, Welch’s t-test). (B) *DCP1A*: siCtrl 51.5 ± 5.2%, siNAT10 35.3 ± 12.6% (p = 0.145; directionally consistent but statistically non-significant decrease, n = 3). (C) *DDX17* and *DCP1A* mRNA enrichment in the Ctrl condition (pcDNA-GFP reporter). HepG2 cells were co-transfected with each FLAG-NAT10 variant and pcDNA-GFP; Anti-FLAG IP and paired IgG IP were performed as described in the Methods (RNA immunoprecipitation section). This condition measures endogenous *DDX17* and *DCP1A* mRNA association with each *NAT10* variant in the absence of overexpressed *DDX17* reporter. Bars represent mean ± SEM (nL=L3). (D) Western blot of FLAG-tagged *NAT10* (NAT10-WT, G641E, and K290A) recovered by anti-FLAG immunoprecipitation from the same lysates used for the RIP-qPCR experiments in Fig.L7C, from cells co-transfected with the DDX17-WT or DDX17-ΔCAG reporter. Comparable band intensity across all six lanes indicates similar immunoprecipitation efficiency of the three *NAT10* variants irrespective of the co-transfected reporter, supporting that the reduced *DDX17* mRNA recovery observed for NAT10-K290A (Fig.L7C) reflects reduced RNA-binding capacity rather than differential immunoprecipitation efficiency.

**Supplementary Figure S8.**
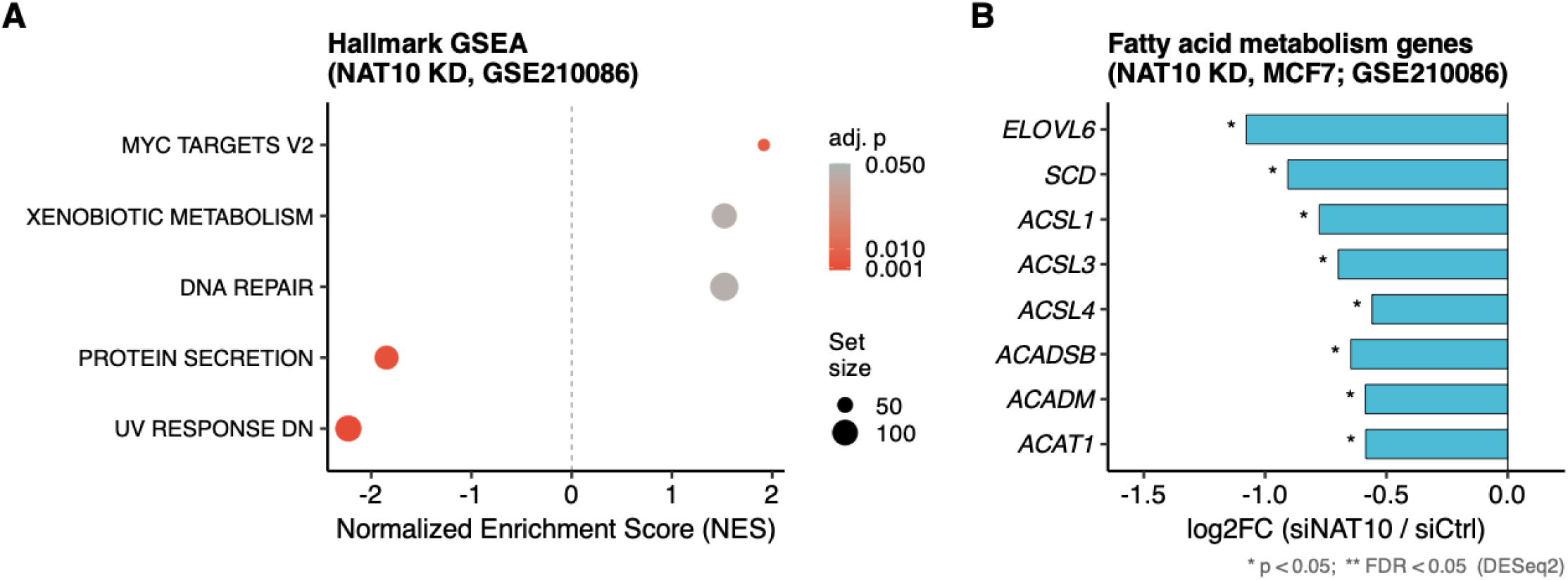
Cross-dataset validation of NAT10-dependent fatty acid metabolic regulation. (A) Gene Set Enrichment Analysis (GSEA) using Hallmark gene sets applied to publicly available *NAT10* knockdown RNA-seq data (GSE210086; siNAT10 vs. siCtrl in MCF7 cells; Dalhat et al., 2022). Significant downregulation of HALLMARK_PROTEIN_SECRETION (NES = −1.85, FDR = 0.002) and HALLMARK_UV_RESPONSE_DN (NES = −2.23, FDR < 0.001) was observed. (B) Differential expression of eight fatty acid metabolism genes identified as NAT10-regulated ac4C targets by Dalhat et al. (ELOVL6, SCD, ACSL1, ACSL3, ACSL4, ACADSB, ACADM, ACAT1) in GSE210086, re-analyzed using DESeq2. All eight genes were consistently downregulated (all p < 0.05), corroborating NAT10-dependent fatty acid metabolic regulation independent of cell type.

**Supplementary Table S3. Convergent evidence supporting NAT10-dependent metabolic and posttranscriptional regulation across independent hepatic and non-hepatic model systems.**

Summary of published studies reporting NAT10-dependent metabolic and posttranscriptional phenotypes in hepatic and non-hepatic model systems. Cell lines or animal models, key endpoints, and alignment with findings in the present study are listed for each independent report. FA, fatty acid; HCC, hepatocellular carcinoma; KD, knockdown; KO, knockout; MASLD, metabolic dysfunction-associated steatotic liver disease; MASH, metabolic dysfunction-associated steatohepatitis.

## STAR METHODS

### KEY RESOURCES TABLE

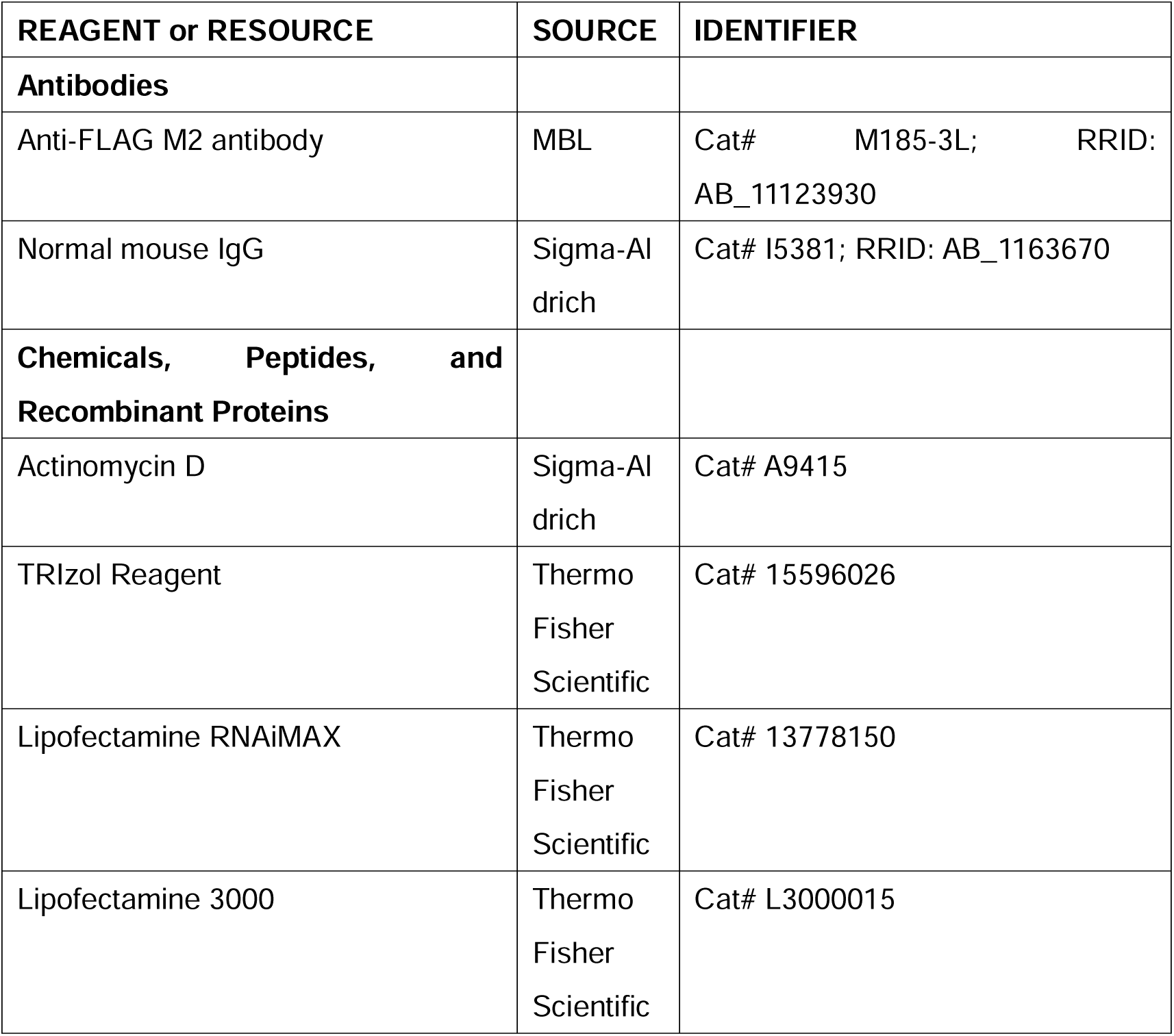
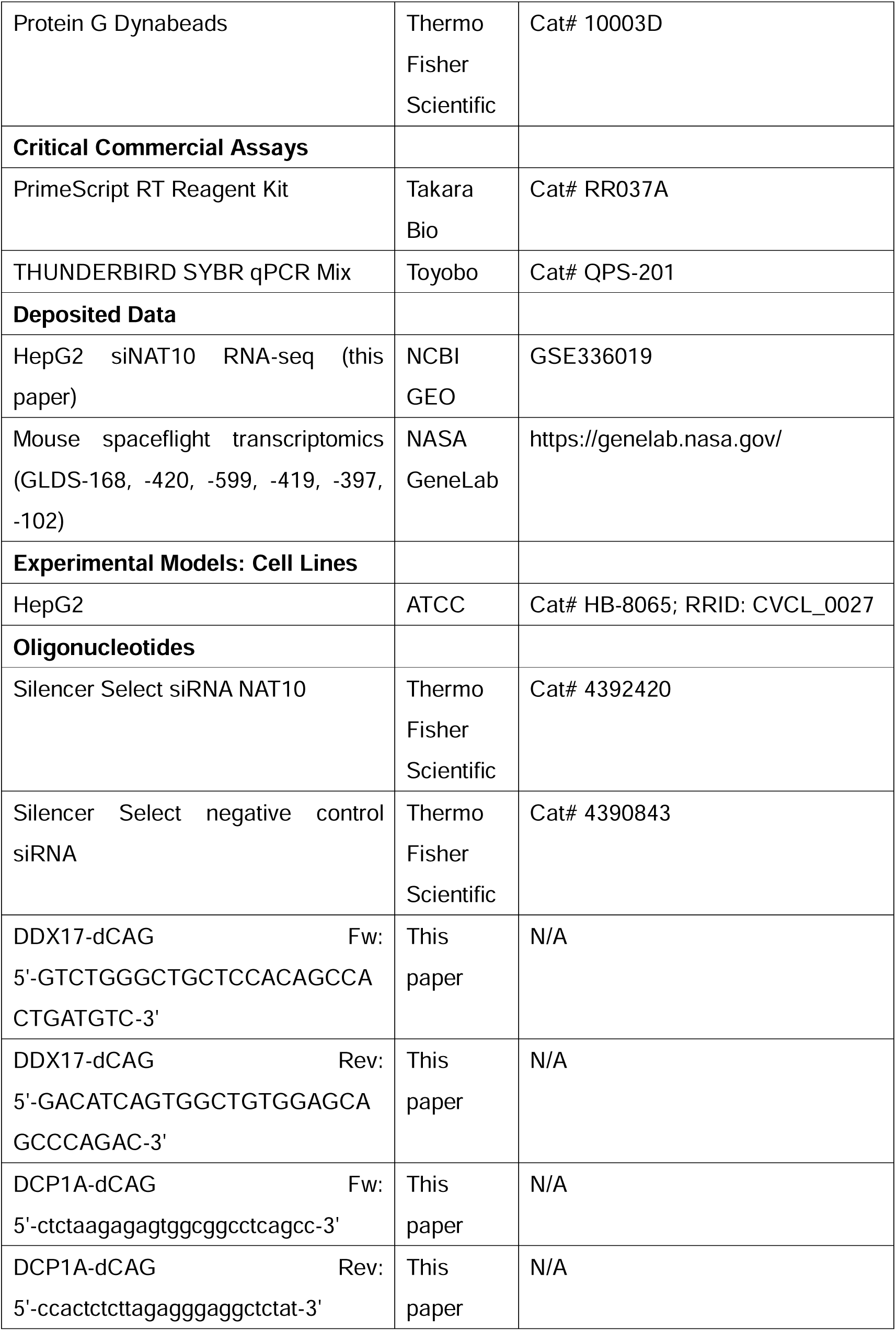
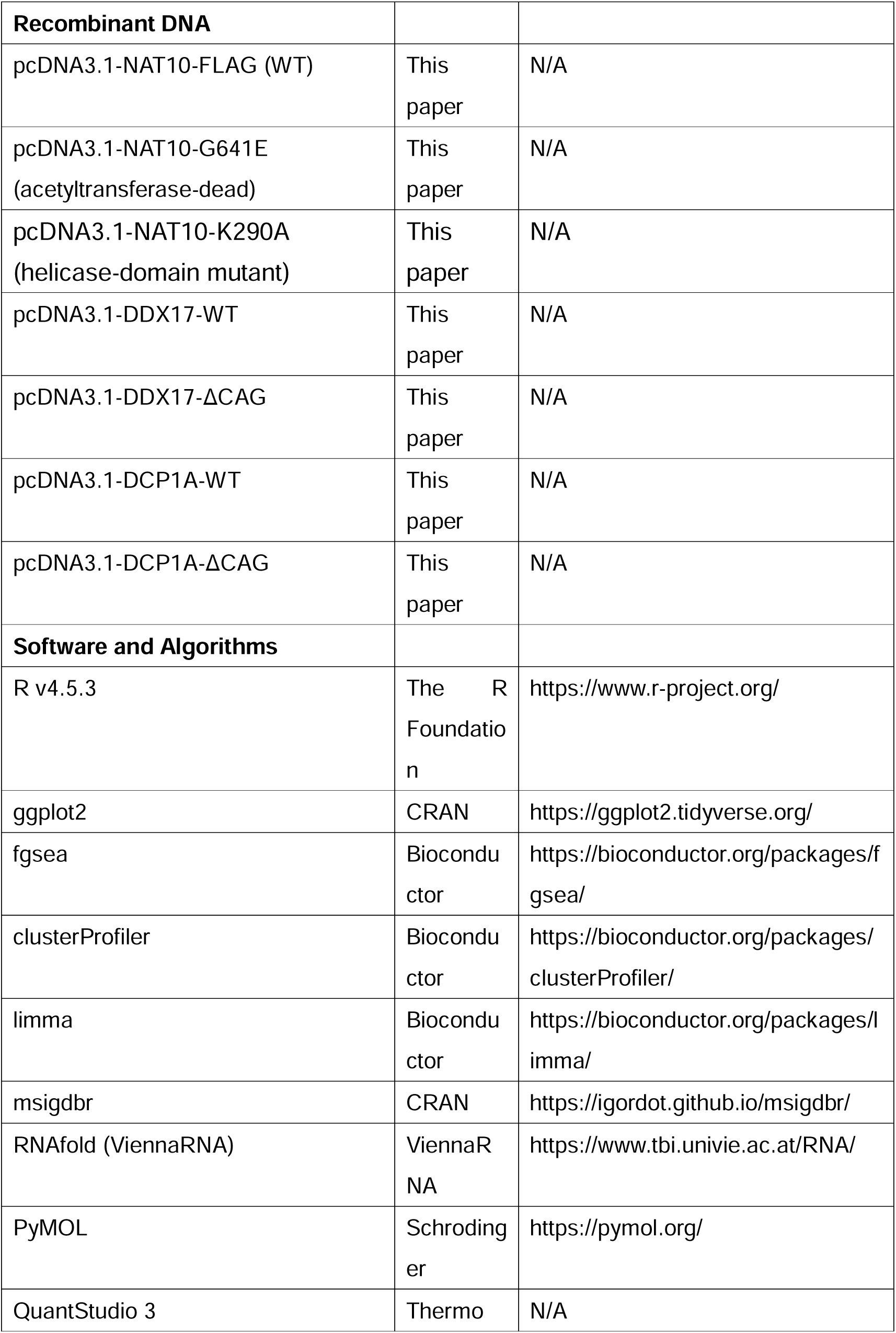
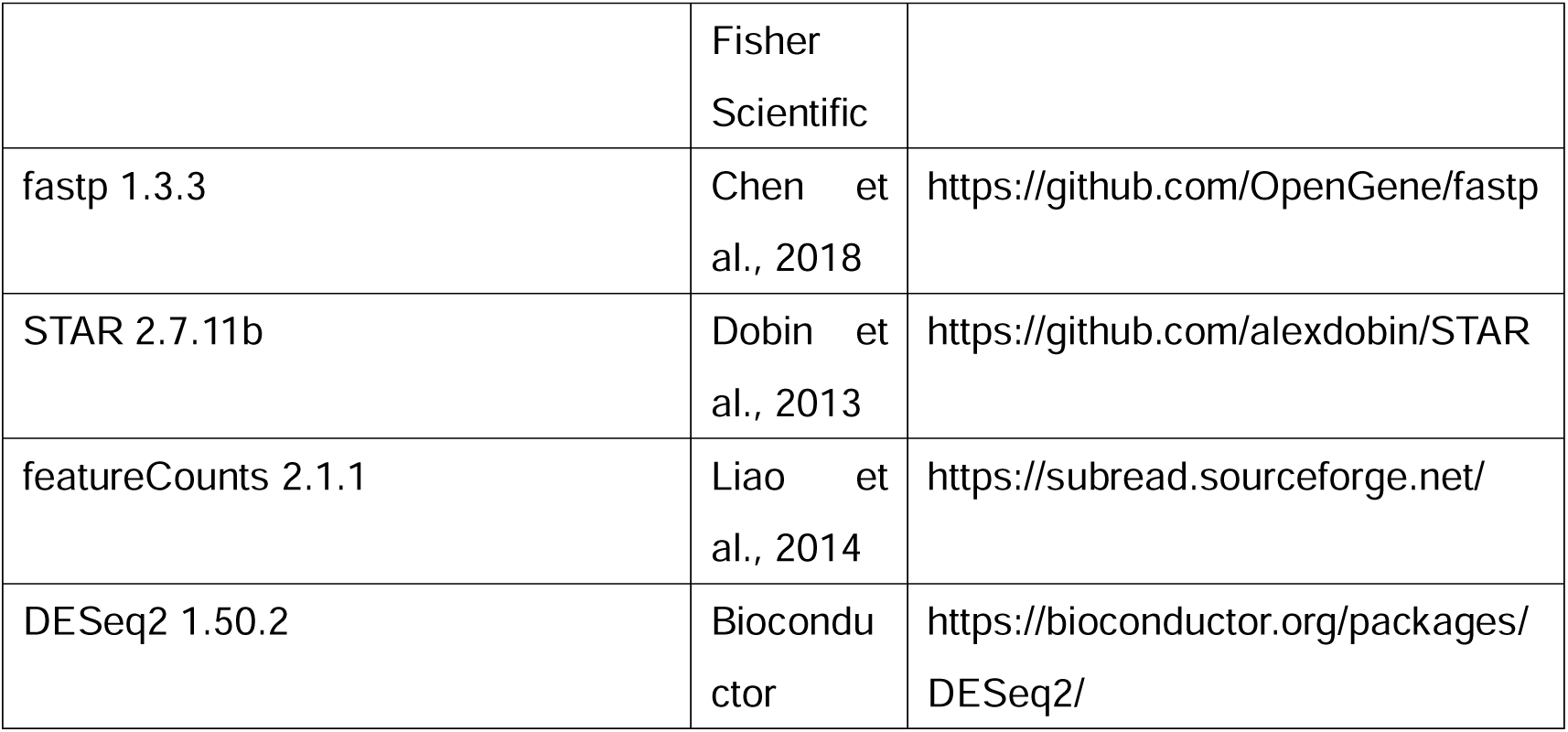

### RESOURCE AVAILABILITY

#### LEAD CONTACT AND MATERIALS AVAILABILITY

Further information and requests for resources and reagents should be directed to and will be fulfilled by the Lead Contact, Ryota Kurimoto.

Plasmids generated in this study are available from the lead contact upon reasonable request.

#### DATA AND CODE AVAILABILITY

All the data required to reproduce this study are included in this published article and supplementary information. The publicly available RNA-seq dataset GSE210086 (Dalhat et al., 2022, NCBI GEO) was re-analyzed in this study; the DESeq2 analysis script is provided as Supplementary Data. Raw data from the qPCR and raw western blot image have been deposited on Mendeley Data, V1, doi: 10.17632/wvfwknrnyx.1. The raw and processed data of RNA-seq (HepG2) generated in this study have been deposited in the Gene Expression Omnibus (GEO) database under accession code GSE336019. Processed expression matrices, sample metadata, motif-analysis source data, and the R scripts used for the analyses are available from the lead contact upon request. Any additional information required to reanalyze the data reported in this paper is available from the lead contact upon request.

## METHOD DETAILS

### EXPERIMENTAL MODEL AND STUDY PARTICIPANT DETAILS

#### Cell culture

HepG2 cells were cultured in Dulbecco’s Modified Eagle’s Medium (DMEM) containing 10% fetal bovine serum (FBS) and 1% penicillin-streptomycin, maintained at 37°C under 5% CO₂.

#### *NAT10* knockdown

*NAT10* was silenced by RNA interference using Silencer Select Pre-Designed siRNA targeting *NAT10* (cat# 4392420, Thermo Fisher Scientific) or a non-targeting control siRNA, delivered with Lipofectamine RNAiMAX (Thermo Fisher Scientific) per the manufacturer’s protocol. Silencing efficacy was verified by RT-qPCR 48 hours after transfection.

#### Plasmid construction and transfection

Expression constructs encoding wild-type *DDX17* (*DDX17* WT) and a single-site CAGCAG-deletion mutant (*DDX17* ΔCAG; one of three CDS CAGCAG hexamers deleted), as well as wild-type *DCP1A* (*DCP1A* WT) and a single-site CAGCAG-deletion mutant (*DCP1A* ΔCAG; one of four CDS CAGCAG hexamers deleted), were cloned into pcDNA3.1 and confirmed by Sanger sequencing. *NAT10* was cloned into pcDNA3.1 using primers Fw: 5’-GAGACCCAAGCTGGCTAGCGTTTAAACTTAAGCTTGGTACCGAGCTCatgcatcggaaaaaggtggataaccgaatccggat-3’, Rev: 5’-AGGCTGATCAGCGGGTTTAAACGGGCCCTCTAGATCAtttcttccgcttcagtttcatatcttttttgttc t-3’, and linearized with BamHI and XhoI (NEB). *NAT10* acetyltransferase-dead (G641E)^13^ plasmid was generated by site-directed mutagenesis using PrimeSTAR Max DNA Polymerase (Takara Bio) with the following primers: Fw: 5’-tgggctatGAGagccgtgctctgcagctg-3’, Rev: 5’-gcacggctCTCatagcccatcccttgata-3’. *NAT10* K290A helicase-domain-mutant plasmid was generated by site-directed mutagenesis using PrimeSTAR Max DNA Polymerase with the following primers: Fw: 5’-gacggggaGCAtctgcagccctgggattg-3’, Rev: 5’-gctgcagaTGCtccccgtcctcgagcagc-3’. Cloning primers for *DDX17* and *DCP1A* were as follows: *DDX17* Fw: 5’-GAGACCCAAGCTGGCTAGCGTTTAAACTTAAGCTTGGTACCGAGCTCATGCCCACCGGCTTTGTAGCCCCGATTCTCTGTGTTTT-3’, *DDX17* Rev: 5’-AGGCTGATCAGCGGGTTTAAACGGGCCCTCTAGATCATCCTCCTCCTCCTCCTGCTGCCTCCTCCGAGCCGGTGTTTACGTGAAGGAGGAGGAGGGGGAGGAGGAGGAG-3’; *DCP1A* Fw: 5’-GAGACCCAAGCTGGCTAGCGTTTAAACTTAAGCTTGGTACCGAGCTCATGGAGGCGCTGAGTCGAGCTGGGCAGGAGATGAG-3’, *DCP1A* Rev: 5’-AGGCTGATCAGCGGGTTTAAACGGGCCCTCTAGATCATCCTCCTCCTCCTCCTG CTGCCTCCTCCGAGCCGGTGTAGGTTGTGGTTGTCTTTGTTCTTGGTCAGAACCT-3’. Single-site CAGCAG deletions were generated by site-directed mutagenesis using PrimeSTAR Max DNA Polymerase (Takara Bio) with the following primers: DDX17-ΔCAG Fw: 5’-GTCTGGGCTGCTCCACAGCCACTGATGTC-3’, Rev: 5’-GACATCAGTGGCTGTGGAGCAGCCCAGAC-3’; DCP1A-ΔCAG Fw: 5’-ctctaagagagtggcggcctcagcc-3’, Rev: 5’-ccactctcttagagggaggctctat-3’. Thus, the ΔCAG constructs retain two of three *DDX17* CAGCAG sites and three of four *DCP1A* CAGCAG sites, respectively. Plasmids were transfected into *NAT10* knockdown cells using Lipofectamine 3000 (Thermo Fisher Scientific) according to the manufacturer’s instructions. Experiments were initiated at 48 hours post-transfection.

#### Actinomycin D mRNA stability assay

mRNA stability was evaluated by actinomycin D (ActD) chase assay. Transcription was blocked by treating cells with actinomycin D (5 μg/mL; A9415, Sigma-Aldrich). For the five-condition rescue assays (Fig.L7A, B), total RNA was extracted at 0, 4, and 8 h post-treatment; for the endogenous siCtrl/siNAT10 assay (Supplementary Fig. LS7A, B), total RNA was extracted at 0 and 4 h post-treatment; RNA was purified using TRIzol (Thermo Fisher Scientific) and quantified by NanoDrop. For domain rescue experiments, HepG2 cells were reverse-transfected with siRNA targeting the 3′UTR of endogenous *NAT10* (siNAT10) or non-targeting control (siCtrl) using Lipofectamine RNAiMAX (Thermo Fisher Scientific). At 24 h post-siRNA transfection, cells were co-transfected with CDS-only rescue plasmids encoding wild-type *NAT10* (pNAT10-WT), acetyltransferase-dead *NAT10* (pNAT10-G641E), or the *NAT10* K290A helicase-domain mutant (pNAT10-K290A); for the siCtrl and siNAT10 conditions, which received no rescue plasmid, pcDNA3.1-GFP was co-transfected in its place to equalize total transfected DNA mass across all five conditions. All conditions were co-transfected with reporter plasmids (*DDX17* and *DCP1A* or DDX17-ΔCAG and DCP1A-ΔCAG), using Lipofectamine 3000. ActD treatment (5 μg/mL) was initiated at 48 h post-siRNA transfection. mRNA remaining at the final timepoint (8 h for the five-condition rescue assays; 4 h for the endogenous assay) was expressed as a percentage relative to the 0 h timepoint within each condition.

#### Reverse transcription and quantitative PCR (RT-qPCR)

Complementary DNA (cDNA) was synthesized from 500 ng total RNA using PrimeScript RT Reagent Kit (Takara Bio) according to the manufacturer’s instructions. Quantitative PCR was performed using THUNDERBIRD SYBR qPCR Mix (Toyobo) on a QuantStudio 3 (Thermo Fisher Scientific) system. For the five-condition rescue reporter assays (Fig.C7A, B), DDX17-WT/DDX17-ΔCAG and DCP1A-WT/DCP1A-ΔCAG reporter transcripts were selectively detected using a common reverse primer annealing within the reporter-appended HiBiT-tag coding sequence (5’-TCATCCTCCTCCTCCTCCTCCGCTG-3’), paired with a gene-specific forward primer within the *DDX17* (5’-CAACAGTTTGCACAGCCTCC-3’) or *DCP1A* (5’-CCAGCTCCCCTTCTCCTCTA-3’) coding sequence; because the reverse primer binds a tag sequence absent from the endogenous transcripts, these primer pairs do not cross-amplify endogenous *DDX17*/*DCP1A* mRNA. For the endogenous ActD assay (Supplementary Fig.CS7A, B), primers targeting the native transcripts were used instead: *DDX17* forward: 5’-ACTGATGCAGCTTGTGGACCAC-3’, reverse: 5’-AAGCCTTCGGTCACACTCATCC-3’; *DCP1A* forward: 5’-cgttgagccctgttctcagt-3’, reverse: 5’-aagatcaacgcttgggaggg-3’. Reference gene primers were: β-actin (ACTB) forward: 5’-CACCATTGGCAATGAGCGGTTC-3’, reverse: 5’-AGGTCTTTGCGGATGTCCACGT-3’; 18S rRNA forward: 5’-CGGCGACGACCCATTCGAAC-3’, reverse: 5’-GAATCGAACCCTGATTCCCCGTC-3’. For all ActD-chase assays (Fig.C7A, B and Supplementary Fig.CS7), *DDX17* and *DCP1A* mRNA levels were normalized to mature 18S rRNA. Although ActD inhibits Pol I-dependent pre-rRNA transcription, mature 18S rRNA is stable over the short assay interval. Relative mRNA remaining was calculated using the comparative Cq (2−ΔΔCq) method as described by Livak and Schmittgen^38^ and expressed relative to the corresponding 0-hour time point within each condition.

#### RNA immunoprecipitation (RIP)

RIP assays were conducted with modifications from a published protocol. For domain-specific binding assays, FLAG-tagged NAT10-WT, NAT10-G641E (acetyltransferase-dead), and NAT10-K290A (helicase-domain mutant) were expressed individually in HepG2 cells and subjected to anti-FLAG immunoprecipitation as described below, followed by RT-qPCR for *DDX17* and *DCP1A* mRNAs.^37^ Cells co-expressing NAT10-FLAG and wild-type or ΔCAG constructs of *DDX17* or *DCP1A* were lysed in RIP buffer (1× PBS, 0.1% SDS, 0.5% sodium deoxycholate, 0.5% NP-40) with rotation at 4°C for 10 min, followed by sonication (30 sec on/30 sec off, 5 cycles). Cleared lysate was obtained by centrifugation (13,000 × g, 15 min, 4°C). Input samples (10% of total lysate) were collected prior to immunoprecipitation; Ct_input values were not mathematically adjusted for this 10% dilution, so the %Input/IP-Input ratio values reported throughout (see formula below) are expressed relative to this 10% input aliquot as collected, rather than extrapolated to 100% of total lysate, and can therefore exceed 1.0 (100%) when IP recovery is high relative to this reference aliquot; this convention is applied identically across all conditions. For immunoprecipitation, mouse anti-FLAG M2 antibody (MBL) or normal mouse IgG (Sigma Aldrich) was coupled to Protein G Dynabeads (Thermo Fisher Scientific) following the manufacturer’s protocol. Coupled antibodies were washed with PBS-T and mixed with whole cell lysate, then rotated at 4°C for 2 hours. Immunocomplexes were captured with a magnetic stand and washed sequentially with low-salt buffer (1× PBS, 0.1% SDS, 0.5% sodium deoxycholate, 0.5% NP-40) and high-salt buffer (5× PBS, same detergents). RNA was recovered by acid phenol-chloroform extraction (Thermo Fisher Scientific). Complementary DNA synthesis and RT-qPCR were performed as described above using the same reporter-specific primers used for the ActD reporter assays (Fig. 7A, B), which selectively detect the co-transfected DDX17-WT/ΔCAG or DCP1A-WT/ΔCAG reporter transcripts. Enrichment was quantified as percent input: % Input = 2^(Ct_input − Ct_IP) × 100. For endogenous target validation, RIP was performed using the same NAT10-FLAG immunoprecipitation protocol described above, without co-transfection of exogenous *DDX17* or *DCP1A* constructs. Enrichment of endogenous *DDX17* and *DCP1A* mRNAs was quantified by RT-qPCR using the endogenous-transcript-targeting primers described above for the endogenous ActD assay (Supplementary Fig. S7A, B), since no reporter construct (and therefore no HiBiT-tag sequence) was present in this condition. Enrichment was expressed as percentage of input (%Input = 2^(Ct_input − Ct_IP) × 100) and as fold enrichment over IgG control.

#### RNA secondary structure prediction and molecular docking

RNA secondary structure of the *DDX17* CAG repeat region (wild-type and single-site ΔCAG sequence) was predicted using RNAfold (ViennaRNA package, v2.7.2). The minimum free energy (MFE) structure and base-pair probability profiles were computed and visualized from the PostScript outputs. The *DDX17* sequence used for structure prediction encompassed 109 nucleotides centered on the selected CAGCAG site deleted in DDX17-ΔCAG; two additional CAGCAG sites elsewhere in the *DDX17* coding sequence remained unchanged in the reporter construct. For molecular docking simulation, three-dimensional all-atom models of the *DDX17* RNA region (wild-type and single-site ΔCAG sequence, 109 nucleotides each) were generated using RNAComposer. The *NAT10* cryo-EM structure (PDB: 9B0E) was docked with a CAG motif RNA sequence using the HDOCK server to establish a reference *NAT10*–RNA complex geometry. Each *DDX17* RNA model was superimposed onto the RNA component of this reference complex using PyMOL (align command), and contact atoms were defined as *NAT10* protein atoms within 5 Å of the superimposed *DDX17* RNA model. Molecular visualization was performed using PyMOL (v3.1.0).

#### Primary liver RNA-seq differential expression analysis (Fig. 1A, B; Supplementary Fig. 5B)

Raw paired-end sequencing reads for the GLDS-168 (OSD-168) mouse liver dataset (FLT, n = 5; GC, n = 5) were downloaded from NASA GeneLab. Reads were quality- and adapter-trimmed using fastp, then aligned to the GENCODE mouse reference genome (GRCm39, primary assembly; GENCODE vM39 annotation) using STAR (v2.7.11b) (--sjdbOverhang 100). Gene-level read counts were quantified using featureCounts (v2.1.1, Subread package); strand specificity was determined empirically on a representative sample and applied uniformly across samples (reverse-stranded protocol, -s 2). Differential expression between FLT and GC was tested using DESeq2 (v1.50.2, Bioconductor) in R (v4.5.3) with the design formula ∼ condition. Low-expression genes were filtered prior to testing, requiring rowSums(counts(dds) ≥ 10) ≥ ceiling(nrow(meta) × 0.2) — i.e., at least 2 of the 10 samples with a count ≥ 10 for a gene to be retained — reducing the gene set from 78,348 to 21,825 genes tested. Significance was assessed using results(dds, alpha = 0.05); no log2 fold-change threshold was imposed in the Wald test itself (null hypothesis: log2 fold change = 0), and the alpha parameter served only to optimize DESeq2’s independent filtering step rather than define the final significance cutoff. padj denotes the Benjamini–Hochberg-adjusted p-value returned by results(); at padj < 0.05, 5,134 genes were significantly differentially expressed overall. For figure display, genes meeting padj < 0.05 were further classified as up- or downregulated using a post hoc |log2 fold change| threshold of > 0.5 (Fig. 1A, genome-wide) or > 0.3 (Fig. 1B, RNA modification enzyme panel; Supplementary Fig. 5B); a more stringent |log2 fold change| ≥ 1 (≥2-fold) threshold was applied separately to define the full significant-gene table (1,489 genes at padj < 0.05 and fold change ≥ 2). *NAT10* was highlighted based on gene symbol matching.

#### Multi-tissue exploratory analysis (**Fig. 1C; Fig. 2–6; Supplementary Figs. 1, 4, 5A**)

For the multi-tissue comparisons, gene coexpression analyses, and the *NAT10* expression boxplot (Fig. 1C), post-normalized TPM values were downloaded directly from NASA GeneLab for six tissues (liver, GLDS-168; spleen, GLDS-420; heart, GLDS-599; muscle, GLDS-419; retina, GLDS-397; kidney, GLDS-102; Table S2) and log2-transformed before analysis. Tissue metadata and sample groupings were derived from the associated GeneLab metadata tables. Because these analyses relied on pre-normalized TPM values rather than raw count matrices, associated statistical comparisons (Welch’s t-test with Benjamini–Hochberg FDR correction, or the Wilcoxon rank-sum test where noted in the figure legends) should be interpreted as screening-level rather than definitive count-based inference. For RNA-seq of NAT10-knockdown HepG2 cells (Fig. 6C), cells were transfected with siRNA targeting *NAT10* or non-targeting control siRNA; total RNA was harvested two days post-transfection. Libraries were prepared using the Lexogen QuantSeq 3′ mRNA-Seq Library Prep Kit FWD and sequenced on an Illumina NovaSeq 6000 (paired-end). Gene expression was quantified using STAR v2.7.9a alignment to GRCh38 followed by featureCounts.

#### Correlation analysis

Spearman’s rank correlation coefficients were computed between *NAT10* and all expressed genes in each tissue, focusing primarily on FLT samples. Genes with high positive correlations (typically top 100–500) were used for motif and GO enrichment analyses. Because FLT-specific rankings were based on five samples, these analyses were treated as exploratory.

#### Gene set enrichment analysis (GSEA)

Differential expression analysis was performed using the limma package (v3.58) with log2-transformed normalized counts. Low-expression genes (log2 expression < 1 in fewer than 3 samples) were excluded. Quantile normalization was applied between arrays. GSEA was performed using the fgsea package (v1.28) with all genes ranked by log2 fold change (FLT vs. GC). Gene sets were retrieved from MSigDB (v2024.1) through the msigdbr R package, including Hallmark gene sets, KEGG Legacy pathways, and Reactome pathways (Mus musculus). Gene set size was restricted to 15–500 terms. Permutations: 1,000. Mouse gene identifiers were mapped to Entrez IDs via the org.Mm.eg.db Bioconductor package. Spearman correlation coefficients between *NAT10* and metabolic pathway activity scores (mean log2 expression of representative pathway genes) were calculated within liver samples (n = 10). As a sensitivity analysis, these correlations were also re-computed within FLT (spaceflight) samples only (n = 5); given the small sample size, exact two-tailed permutation p-values (cor.test(…, method = "spearman", exact = TRUE) in R) were used rather than the asymptotic t-approximation. All computational analyses were conducted using R (v4.5.3).

#### Motif scanning and positional mapping

Coding sequences (CDS) and 3′UTR regions were extracted from Ensembl FASTA files (GRCm39) using transcript mapping and annotation parsing. Sequence scanning for motifs (e.g., CAGCAG, CXXC, ARE, PRE, CU-rich, IGF2BP targets) was performed using Biostrings with regular expression matching. Each motif was assigned a genomic region (CDS or UTR) for stratified analysis.

#### Motif enrichment testing

Motif enrichment among *NAT10*-correlated gene sets was assessed by constructing 2×2 contingency tables comparing motif-positive and motif-negative genes within the top N correlated genes versus all other genes. Fisher’s exact test was used to estimate odds ratios and p-values, and plotted for selected motifs. To generate a background set for motif enrichment analysis, we randomly sampled 500 coding sequences (CDS) from the Mus musculus GRCm39 reference transcriptome (Ensembl release). Random sampling was performed with a fixed seed for reproducibility, and the resulting set (Random500_CDS.fasta) was used for background normalization of motif frequencies. The matched-background empirical tests in Fig. 3J and K were considered the primary sensitivity analyses; nominal Fisher p-values from the broader motif screen were treated as exploratory.

#### Gene Ontology (GO) and GSEA analyses

GO enrichment was performed using clusterProfiler and msigdbr packages with background defined as all expressed genes. Gene Set Enrichment Analysis (GSEA) was applied using preranked lists sorted by *NAT10* correlation or logFC. Pathways related to RNA processing, splicing, and decay were specifically evaluated.

#### RNA modification gene profiling

A manually curated list of RNA modification-associated genes (writers, readers, erasers) was assembled and analyzed for logFC and expression trends across FLT and GC groups. PCA was conducted on variance-stabilized expression profiles. For PCA visualization, a representative subset of major RNA modification-related genes was used.

### QUANTIFICATION AND STATISTICAL ANALYSIS

Data visualization was performed using ggplot2, pheatmap, and enrichplot. Analyses were conducted in R (v4.5.3) with Bioconductor packages. Multiple-testing correction was performed using the Benjamini–Hochberg method unless otherwise noted. Statistical tests were selected according to the experimental design and are specified in the corresponding Methods subsections and figure legends. Unless otherwise indicated, bar graphs show mean ± SEM; Supplementary Fig. S7A and B show mean ± SD; box plots show the median and interquartile range. Statistical significance was set at p < 0.05. All in vitro assays were performed in three independent biological experiments (n = 3), with consistent directional results across experiments.

